# Neutralization of SARS-CoV-2 by destruction of the prefusion Spike

**DOI:** 10.1101/2020.05.05.079202

**Authors:** Jiandong Huo, Yuguang Zhao, Jingshan Ren, Daming Zhou, Helen ME Duyvesteyn, Helen M Ginn, Loic Carrique, Tomas Malinauskas, Reinis R Ruza, Pranav NM Shah, Tiong Kit Tan, Pramila Rijal, Naomi Coombes, Kevin Bewley, Julika Radecke, Neil G Paterson, Piyasa Supasa, Juthathip Mongkolsapaya, Gavin R Screaton, Miles Carroll, Alain Townsend, Elizabeth E Fry, Raymond J Owens, David I Stuart

**Author notes:** Correspondence: David I. Stuart (Lead Contact), or Jingshan Ren. These authors contributed equally to this work (Jiandong Huo, Yuguang Zhao).

## Abstract

There are as yet no licenced therapeutics for the COVID-19 pandemic. The causal coronavirus (SARS-CoV-2) binds host cells via a trimeric Spike whose receptor binding domain (RBD) recognizes angiotensin-converting enzyme 2 (ACE2), initiating conformational changes that drive membrane fusion. We find that monoclonal antibody CR3022 binds the RBD tightly, neutralising SARS-CoV-2 and report the crystal structure at 2.4 Å of the Fab/RBD complex. Some crystals are suitable for screening for entry-blocking inhibitors. The highly conserved, structure-stabilising, CR3022 epitope is inaccessible in the prefusion Spike, suggesting that CR3022 binding would facilitate conversion to the fusion-incompetent post-fusion state. Cryo-EM analysis confirms that incubation of Spike with CR3022 Fab leads to destruction of the prefusion trimer. Presentation of this cryptic epitope in an RBD-based vaccine might advantageously focus immune responses. Binders at this epitope may be useful therapeutically, possibly in synergy with an antibody blocking receptor attachment.

**Highlights:**

- **CR3022 neutralises SARS-CoV-2**
- **Neutralisation is by destroying the prefusion SPIKE conformation**
- **This antibody may have therapeutic potential alone or with one blocking receptor attachment**

## Introduction

Incursion of animal (usually bat)-derived coronaviruses into the human population has caused several outbreaks of severe disease, starting with severe acute respiratory syndrome (SARS) in 2002 (Menachery et al., 2015). In late 2019 a highly infectious illness, with cold-like symptoms progressing to pneumonia and acute respiratory failure, resulting in an estimated 6% overall death rate (Baud et al., 2020), with higher mortality among the elderly and immunocompromised populations, was identified and confirmed as a pandemic by the WHO on 11^th^ March 2020. The etiological agent is a novel coronavirus (SARS-CoV-2) belonging to lineage B betacoronavirus and sharing 88% sequence identity with bat coronaviruses (Lu et al., 2020a). The heavily glycosylated trimeric surface Spike protein mediates viral entry into the host cell. It is a large type I transmembrane glycoprotein (the ectodomain alone comprises over 1200 residues) (Wrapp et al., 2020). It is made as a single polypeptide and then cleaved by host proteases to yield an N-terminal S1 region and the C-terminal S2 region. Spike exists initially in a pre-fusion state where the domains of S1 cloak the upper portion of the spike with the relatively small (∼22 kDa) S1 RBD nestled at the tip. The RBD is predominantly in a ‘down’ state where the receptor binding site is inaccessible, however it appears that it stochastically flips up with a hinge-like motion transiently presenting the ACE2 receptor binding site (Roy, 2020; Song et al., 2018; Walls et al., 2020; Wrapp et al., 2020). ACE2 acts as a functional receptor for both SARS-CoV and SARS-CoV-2, binding to the latter with a 10 to 20-fold higher affinity (K_D_ of ∼15 nM), possibly contributing to its ease of transmission (Song et al., 2018; Wrapp et al., 2020). There is 73% sequence identity between the RBDs of SARS-CoV and SARS-CoV-2 (Figure S1). When ACE2 locks on it holds the RBD ‘up’, destabilising the S1 cloak and possibly favouring conversion to a post-fusion form where the S2 subunit, through massive conformational changes, propels its fusion domain upwards to engage with the host membrane, casting off S1 in the process (Song et al., 2018; Wrapp et al., 2020). Structural studies of the RBD in complex with ACE2 (Lan et al., 2020; Wang et al., 2020b; Yan et al., 2020) how that it is recognized by the extracellular peptidase domain (PD) of ACE2 through mainly polar interactions. The S protein is an attractive candidate for both vaccine development and immunotherapy. Potent nanomolar affinity neutralising human monoclonal antibodies against the SARS-CoV RBD have been identified that attach at the ACE2 receptor binding site (including M396, CR3014 and 80R (Ter Meulen et al., 2006; Sui et al., 2004; Zhu et al., 2007)). For example 80R binds with nanomolar affinity, prevents binding to ACE2 and the formation of syncytia *in vitro*, and inhibits viral replication *in vivo* (Sui et al., 2004). However, despite the two viruses sharing the same ACE2 receptor these ACE2 blocking antibodies do not bind SARS-CoV-2 RBD (Wrapp et al., 2020). In contrast CR3022, a SARS-CoV-specific monoclonal selected from a single chain Fv phage display library constructed from lymphocytes of a convalescent SARS patient and reconstructed into IgG1 format (Ter Meulen et al., 2006), has been reported to cross-react strongly, binding to the RBD of SARS-CoV-2 with a K_D_ of 6.3 nM (Tian et al., 2020), whilst not competing with the binding of ACE2 (Ter Meulen et al., 2006). Furthermore, although SARS-CoV escape mutations could be readily generated for ACE2-blocking CR3014, no escape mutations could be generated for CR3022, preventing mapping of its epitope (Ter Meulen et al., 2006). Furthermore a natural mutation of SARS-CoV-2 has now been detected at residue 495 (Y→N) (GISAID (Shu and McCauley, 2017): Accession ID: EPI_ISL_429783 Wienecke-Baldacchino et al., 2020), which forms part of the ACE2 binding epitope. Finally, CR3022 and CR3014 act synergistically to neutralise SARS-CoV with extreme potency (Ter Meulen et al., 2006). Whilst this work was being prepared for publication a paper reporting that CR3022 does not neutralise SARS-CoV-2 and describing the structure of the complex with the RBD at 3.1 Å resolution was published (Yuan et al., 2020). Here we extend the structure analysis to significantly higher resolution and, using a different neutralisation assay, show that CR3022 does neutralise SARS-CoV-2, but via a mechanism that would not be detected by the method of Yuan *et al* (Yuan et al., 2020). We use cryo-EM analysis of the interaction of CR3022 with the full Spike ectodomain to confirm this mechanism. Taken together these observations suggest that the CR3022 epitope should be a major target for therapeutic antibodies.

## Results

### CR3022 binds tightly to the RBD and allosterically perturbs ACE2 binding

To understand how CR3022 works we first investigated the interaction of CR3022 Fab with isolated recombinant SARS-CoV-2 RBD, both alone and in the presence of ACE2. Surface plasmon resonance (SPR) measurements (Methods and Figure S2) confirmed that CR3022 binding to RBD is strong (although weaker than the binding reported to SARS-CoV (Ter Meulen et al., 2006)), with a slight variation according to whether CR3022 or RBD is used as the analyte (K_D_ = 30 nM and 15 nM respectively, derived from the kinetic data in Table S1). An independent measure using Bio-Layer Interferometry (BLI) with RBD as analyte gave a K_D_ of 19 nM (Methods and Figure S2). These values are quite similar to those reported by Tian *et al*. (Tian et al., 2020) (6.6 nM), whereas weaker binding (K_D_ ∼ 115 nM) was reported recently by Yuan *et al*. (Yuan et al., 2020). Using SPR to perform a competition assay revealed that the binding of ACE2 to the RBD is perturbed by the presence of CR3022 (Figure S3). The presence of ACE2 slows the binding of CR3022 to RBD and accelerates the dissociation. Similarly, the release of ACE2 from RBD is accelerated by the presence of CR3022. These observations are suggestive of an allosteric effect between ACE2 and CR3022.

### CR3022 neutralises SARS-CoV-2

A plaque reduction neutralisation test using SARS-CoV-2 virus and CR3022 showed an ND50 of 1:201 for a starting concentration of 2mg/mL (calculated according to Grist (Grist, 1966)), superior to that of MERS convalescent serum (ND50 of 1:149) used as a NIBSC international standard positive control (see Methods and Table S2). This corresponds to 50% neutralisation at ∼70 nM (∼10.5 ug/mL). This is similar to the neutralising concentration (50% neutralisation at 11 ug/mL) reported by Ter Meulen *et al*. (Ter Meulen et al., 2006) for SARS-CoV, however, as discussed below, it is in apparent disagreement with the result reported recently by Yuan *et al*. (Yuan et al., 2020).

### Structure determination of RBD-CR3022 Fab complex

We determined the crystal structure of the SARS-CoV-2 RBD-CR3022 Fab complex (see Methods and Table S3) to investigate the relationship between the binding epitopes of ACE2 and CR3022. Crystals grew rapidly and consistently. Two crystal forms grew in the same drop. The solvent content of the crystal form solved first was unusually high (ca 87%) with the ACE2 binding site exposed to large continuous solvent channels within the crystal lattice (Figure S4). These crystals therefore offer a promising vehicle for crystallographic screening to identify potential therapeutics that could act to block virus attachment. The current analysis of this crystal form is at 4.4 Å resolution and so, to avoid overfitting, refinement used a novel real-space refinement algorithm to optimise the phases (Vagabond, HMG unpublished, see Methods). This, together with the favourable observation to parameter ratio resulting from the exceptionally high solvent content, meant that the map was of very high quality, allowing reliable structural interpretation (Figure S5, Methods). Full interpretation of the detailed interactions between CR3022 and the RBD was enabled by the second crystal form which diffracted to high resolution, 2.4 Å, and the structure of which was refined to give an R-work/R-free of 0.213/0.239 and good stereochemistry (Methods, Table S3, Figure S5).

### CR3022 binding epitope is highly conserved and inaccessible in prefusion S protein

The high-resolution structure is shown in Figure 1a. There are two complexes in the crystal asymmetric unit with residues 331-529 in one RBD, 332-445 and 448-532 in the other RBD well defined, whilst residues133-136 of the CR3022 heavy chains are disordered. The RBD has a very similar structure to that seen in the complex of SARS-CoV-2 RBD with ACE2, rmsd for 194 Ca atoms of 0.6 Å^2^ (PDB, 6M0J (Lan et al., 2020)), and an rmsd of 1.1 Å^2^ compared to the SARS CoV RBD (PDB, 2AJF (Li et al., 2005)). Only minor conformational changes are introduced by binding to CR3022, at residues 381-390. The RBD was deglycosylated (Methods) to leave a single saccharide unit at each of the N-linked glycosylation sites clearly seen at N331 and N343 (Figure S5). CR3022 attaches to the RBD surface orthogonal to the ACE2 receptor binding site. There is no overlap between the epitopes and indeed both the Fab and ACE2 ectodomain can bind without clashing (Figure 1d) (Tian et al., 2020). Such independence of the ACE2 binding site has been reported recently for another SARS-CoV-2 neutralising antibody, 47D11 (Wang et al., 2020a). The Fab complex interface buries 990 Å^2^ of surface area (600 and 390 Å^2^ by the heavy and light chains respectively, Figure 2a and Figure S6), somewhat more than the RBD-ACE2 interface which covers 850 Å^2^ (PDB 6M0J (Lan et al., 2020)). Typical of a Fab complex, the interaction is mediated by the antibody CDR loops, which fit well into the rather sculpted surface of the RBD (Figure 1b, c). The heavy chain CDR1, 2 and 3 make contacts to residues from α2, β2 and α3 (residues 369-386), while two of the light chain CDRs (1 and 2) interact mainly with residues from the β2-α3 loop, α3 (380-392) and the α5-β4 loop (427-430) (Figures 1, S1, S7). A total of 16 residues from the heavy chain and 14 from the light chain cement the interaction with 26 residues from the RBD. For the heavy chain these potentially form 7 H-bonds and 3 salt bridges, the latter from D55 and E57 (CDR2) to K378 of the RBD. Whilst the light chain interface comprises 6 H-bonds and a single salt bridge between E61 (CDR2) and K386 of the RBD. The binding is consolidated by a number of hydrophobic interactions (Figure S7b). Of the 26 residues involved in the interaction 23 are conserved between SARS-CoV and SARS-CoV-2 (Figure 2b and Figure S1). The CR30222 epitope is much more conserved than that of the receptor blocking anti-SARS-CoV antibody 80R for which only 13 of the 29 interacting residues are conserved (Hwang et al., 2006), in-line with the lack of cross reactivity observed for the latter.

**Figure 1.**
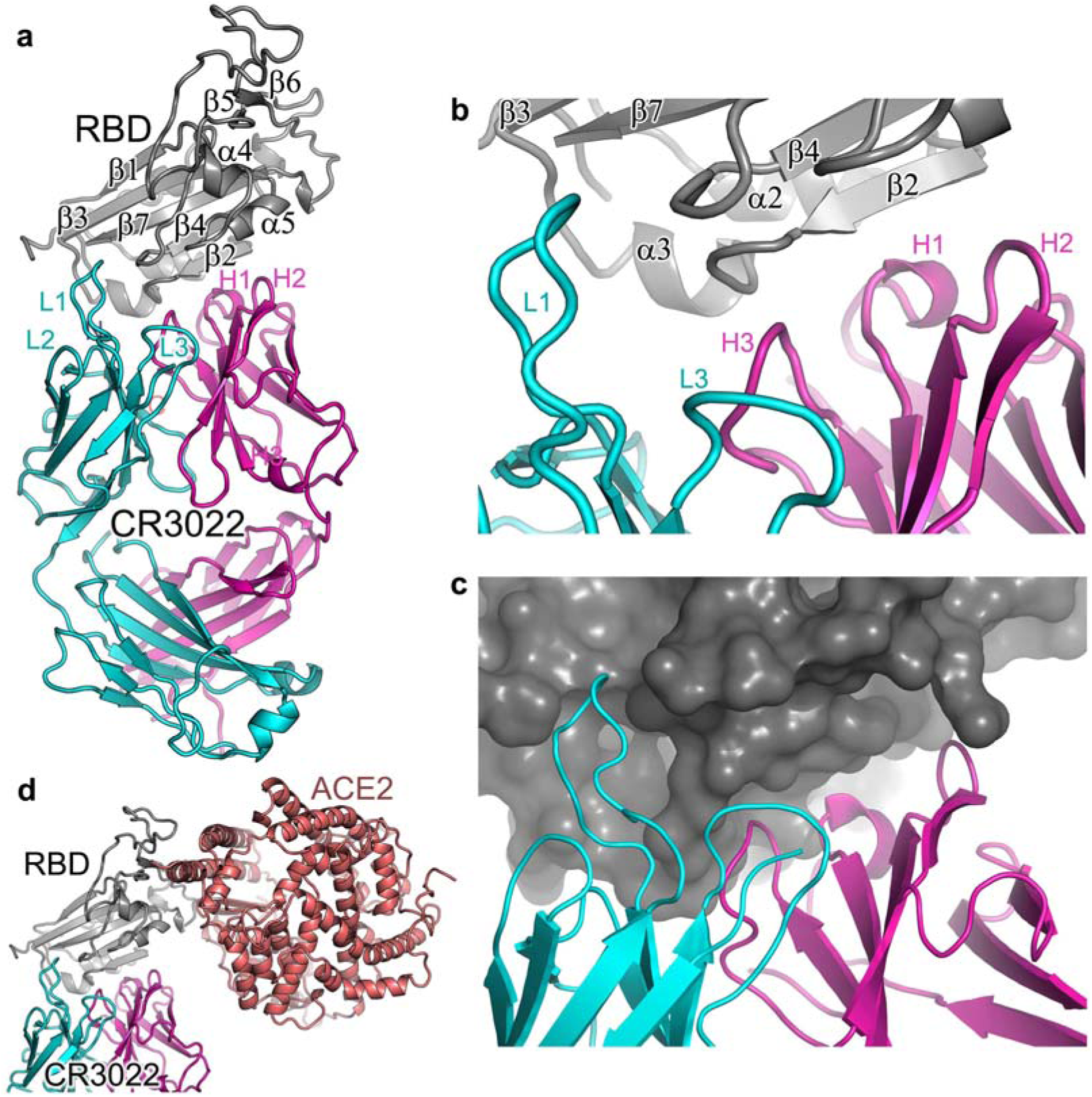
Overall structure of RBD/CR3022 complex. **a**, Ribbon diagram showing the structure of the RBD/CR3022 complex with the RBD shown in grey, CR3022 heavy chain in magenta and light chain in cyan. The heavy chain CDR1-3 are labelled as H1-H3 and the light chain CDR1-3 as L1-3 (where visible). **b**, Closeup of the antigen-antibody binding interface in cartoon representation. **c**, similar view to (**b**) but showing the RBD as a surface. **d**, The RBD of the RBD/ACE2 complex has been overlapped with the RBD of the RBD/CR3022 complex to show the relative positions of the antigenic and receptor binding sites. ACE2 is drawn as a salmon ribbon.

**Figure 2.**
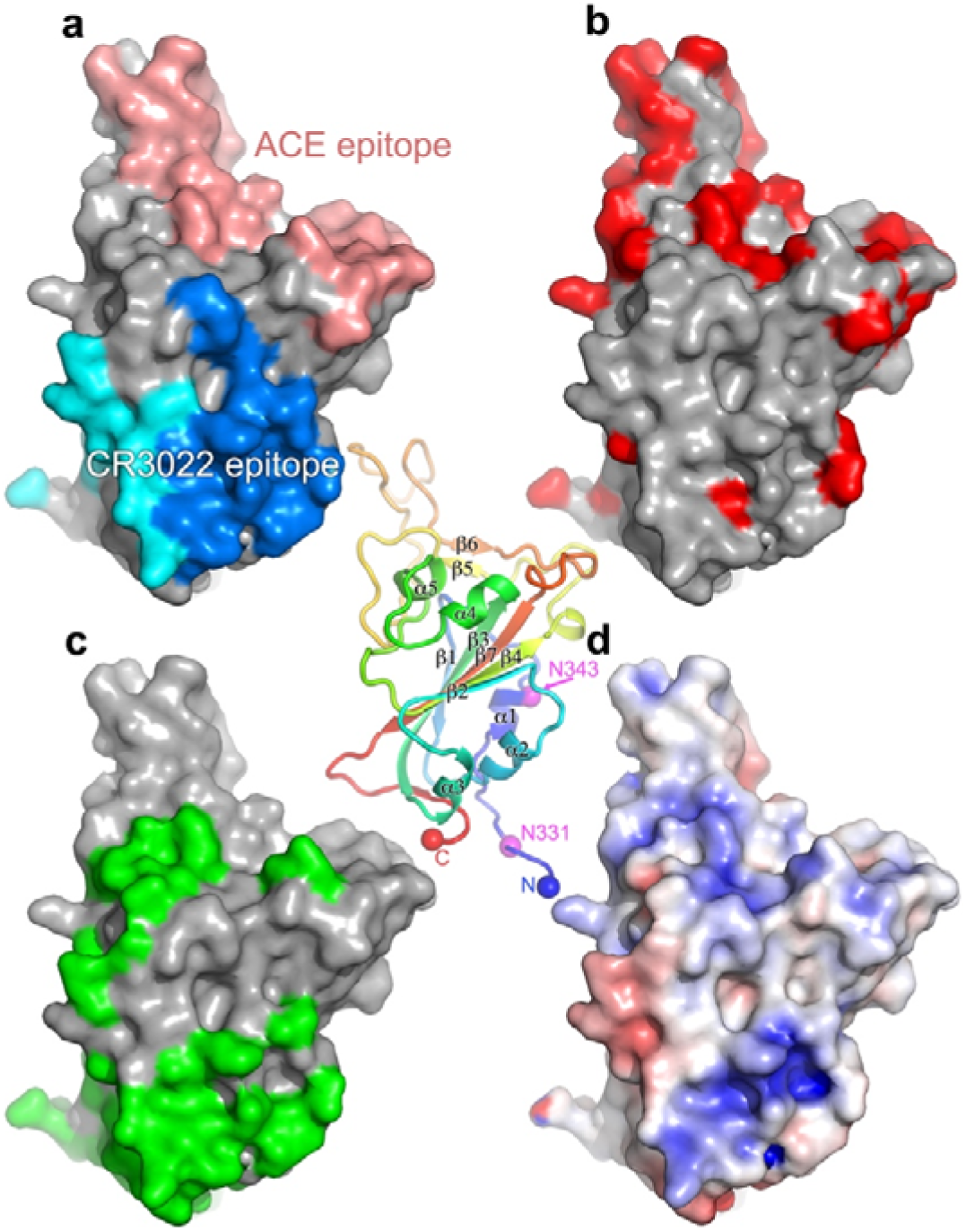
Surface properties of SARS-CoV-2 RBD. The central panel is a cartoon depiction rainbow coloured from blue for the N-terminus to red for the C-terminus, the view is the same as for panels a-d. The secondary structure is labelled along with the glycosylated residue N343 (in magenta) and the position of the domain termini (N and C). **a**, Surface representation of RBD with the solvent accessible area buried by ACE2 receptor binding coloured in salmon, and that buried by CR3022 (heavy chain in blue and light chain in cyan). **b**, Sequence differences shown in red between SARS-CoV and SARS-CoV-2 RBDs, mapped on the surface of SARS-CoV-2 RBD. **c**, The surface buried in the pre-fusion conformation of the Spike shown in green. **d**, The electrostatic surface of SARS-CoV-2 RBD contoured at ± 5 T/e (red, negative; blue, positive).

The reason for the conservation of the CR3022 epitope becomes clear in the context of the complete pre-fusion S structure (PDB IDs: 6VSB (Wrapp et al., 2020), 6VXX, 6VYB (Walls et al., 2020)) where the epitope is inaccessible (Figure 3). When the RBD is in the ‘down’ configuration the CR3022 epitope is packed tightly against another RBD of the trimer and the N-terminal domain (NTD) of the neighbouring protomer. In the structure of the pre-fusion form of trimeric Spike the majority of RBDs are ‘down’, although presumably stochastically one may be ‘up’ (Walls et al., 2020; Wrapp et al., 2020). The structure of a SARS-CoV complex with ACE2 ectodomain shows that this ‘up’ configuration is competent to bind receptor, and that there are a family of ‘up’ orientations with significantly different hinge angles (Song et al., 2018). However, the CR3022 epitope remains largely inaccessible even in the ‘up’ configuration. Modelling the rotation of the RBD required to enable Fab interaction in the context of the Spike trimer, showed a rotation corresponding to a > 60° further declination from the central vertical axis was required, beyond that observed previously (Walls et al., 2020; Wrapp et al., 2020) (Figure 3i), although this might be partly mitigated by more complex movements of the RBD and if more than one RBD is in the ‘up’ configuration this requirement would be relaxed somewhat. Since locking the up state by receptor blocking antibodies is thought to destabilise the pre-fusion state (Walls et al., 2019) binding of CR3022 presumably introduces further destabilisation, leading to a premature conversion to the post-fusion state, inactivating the virus. CR3022 and ACE2 blocking antibodies can bind independently but both induce an ‘up’ conformation, presumably explaining the observed synergy between binding at the two sites (Ter Meulen et al., 2006).

**Figure 3.**
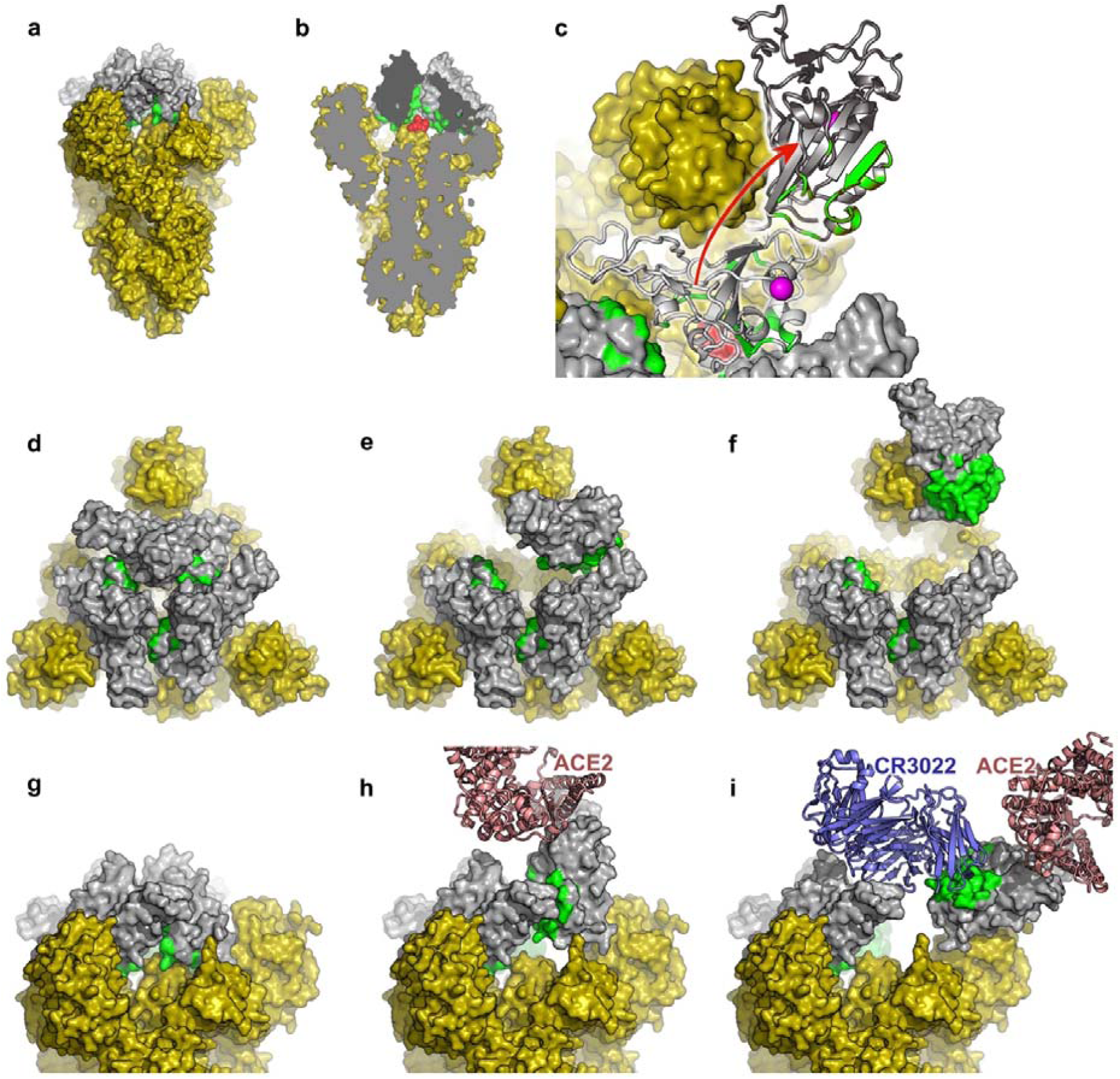
The CR3022 binding regions are inaccessible in the pre-fusion form of the S protein. Panels **a-c** provide an overview. **a**, The pre-fusion state of the S protein with all RBDs in the down conformation (generated by superposing our RBD structure on the pre-fusion trimer of ref (Wrapp et al., 2020)). The viral membrane would be at the bottom of the picture. All of S1 and S2 are shown in yellow, apart from the RBD, which is shown in grey, with the CR3022 epitope coloured green. **a**, A cut-way of the trimer showing, in red, the di-peptide (residues 986-987) which has been mutated to PP to confer stability on the pre-fusion state. Note the proximity to the CR3022 epitope. **c**, Showing a top view of the molecule (also used for panels **d-f**). One of the RBDs has been drawn in light grey in the down configuration and hinged up in dark grey, using the motion about the hinge axis observed for several coronavirus Spikes, but extending the motion sufficiently to allow CR3022 to bind. The PP motif is shown in red and the glycosylated residue N343 in magenta. Panels **d-f** show the trimer viewed from above **d** – all RBDs down, **e** – one RBD up **f** – one RBD rotated (as in **c**) to allow access to CR3022. Panels **g-i** are equivalent structures to **d-f**, but are viewed from the side. In **e** bound ACE2 is shown and in **f** CR3022.

### Mechanism of neutralisation of SARS-CoV-2 by CR3022 confirmed by cryo-EM

To test if CR3022 binding destabilises the prefusion state of Spike, the ectodomain construct described previously (Wrapp et al., 2020) was used to produce glycosylated protein in HEK cells (Methods). Cryo-EM screening showed that the protein was in the trimeric prefusion conformation. Spike was then mixed with an excess of CR3022 Fab and incubated at room temperature, with aliquots being taken at 50 minutes and 3 hours. Aliquots were immediately applied to cryo-EM grids and frozen (Methods). For the 50 minutes incubation, collection of a substantial amount of data allowed unbiased particle picking and 2D classification which revealed two major structural classes with a similar number in each, (i) the prefusion conformation, and (ii) a radically different conformation (Methods, Table S4 and Figure S8). Detailed analysis of the prefusion conformation led to a structure at a nominal resolution of 3.4 Å (FSC = 0.143), based on a broad distribution of orientations, that revealed the same predominant RBD pattern (one ‘up’ and two ‘down’) previously seen (Wrapp et al., 2020) with no evidence of CR3022 binding (Figure 4a, Figure S9). Analysis of the other major particle class revealed strong preferential orientation of the particles on the grid (Figure S10a). Despite this a reconstruction with a nominal resolution of 3.9 Å within the plane of the grid, and perhaps 7 Å resolution in the perpendicular direction (Figure S10b), could be produced which allowed the unambiguous fitting of the CR3022-RBD complex (Figure 4b). Note that in addition there is less well defined density attached to the RBD, in a suitable position to correspond to the Spike N-terminal domain (Wrapp et al., 2020). These structures are no longer trimeric, rather two complexes associate to form an approximately symmetric dimer (however, application of this symmetry in the reconstruction process did not improve the resolution). The interactions responsible for dimerisation involve the ACE2 binding site on the RBD and the elbow of the Fab, however the interaction does not occur in our low-resolution crystal form and is therefore probably extremely weak and not biologically significant. Since conversion to the post-fusion conformation leads to dissociation of S1 (which includes the N-terminal domain and RBD) these results confirm that CR3022 destabilises the prefusion Spike conformation. Further evidence of this is provided by analysis of data collected after 3 h incubation. By this point there were no intact trimers remaining and a heterogeneous range of oligomeric assemblies had appeared, which we were not able to interpret in detail but which are consistent with the lateral assembly of Fab/RBD complexes (Figure S11). Note that the relatively slow kinetics will not be representative of events *in vivo*, where the conversion might be accelerated by the elevated temperature and the absence of the mutations which were added to this construct to stabilise the prefusion state (Kirchdoerfer et al., 2018; Pallesen et al., 2017; Wrapp et al., 2020).

**Figure 4.**
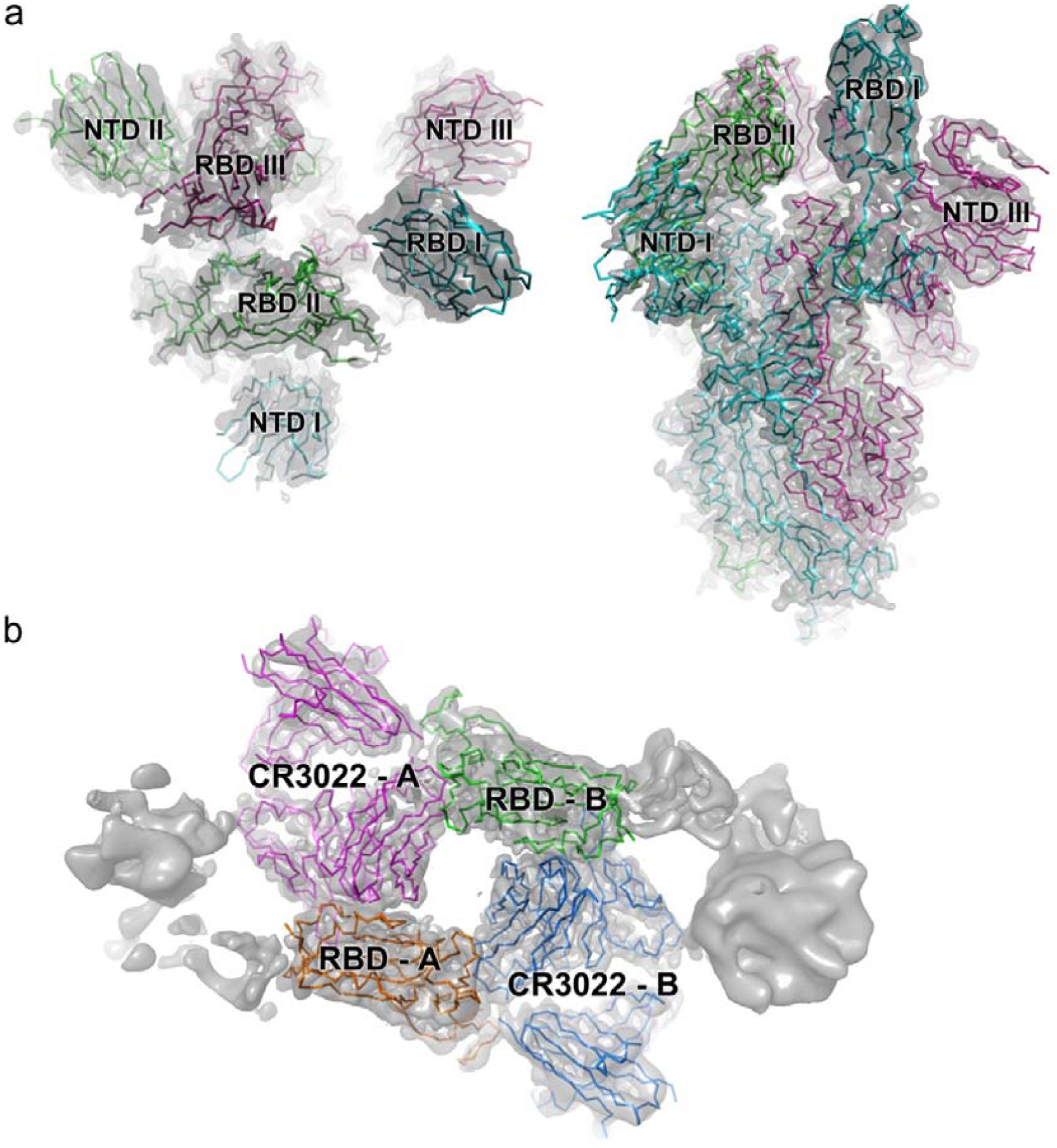
Cryo-EM reconstructions. **a**, shows the prefusion Spike, left top view, right side view. Note RBD I is in the ‘up’ conformation. **b**, shows the dimeric RBD/CR3022 complex, the two complexes are labelled A and B.

## Discussion

Until now the only documented mechanism of neutralisation of coronaviruses has been through blocking receptor attachment. In the case of SARS-CoV this is achieved by presentation of the RBD of the Spike in an ‘up’ conformation. Although not yet confirmed for SARS-CoV-2 it is very likely that a similar mechanism can apply. Here we define a second class of neutralisers, that bind a highly conserved epitope (Figure S1) and can therefore act against both SARS-CoV and SARS-CoV-2 (CR3022 was first identified as a neutralising antibody against SARS-CoV (Ter Meulen et al., 2006)). We find that binding of CR3022 to the isolated RBD is tight (∼20 nM) and the crystal structure of the complex reveals the atomic detail of the interaction. Despite the spatial separation of the CR3022 and ACE2 epitopes we find an allosteric effect between the two binding events. The role of the CR3022 epitope in stabilising the prefusion Spike trimer explains why it has, to date, proved impossible to generate mutations that escape binding of the antibody (Ter Meulen et al., 2006).

Whilst in our assay CR3022 neutralises SARS-CoV-2, a recent paper (Yuan et al., 2020) reported an alternative assay that did not detect neutralisation. The difference is likely due to their removal of the antibody/virus mix after adsorption to the indicator cells, before incubating to allow cytopathic effect (CPE) to develop. This would be in-line with the distinction previously seen between neutralisation tests for influenza virus by antibodies which bind the stem of hemagglutinin and therefore do not block receptor binding (Thomson et al., 2012). These antibodies did not appear to be neutralising when tested with the standard WHO neutralisation assay, in which a similar protocol is used to that adopted by Yuan et al, in which the inoculum of virus/antibody is washed out before development of CPE. Neutralisation was observed, however, when the antibodies were left in the assay during incubation to produce CPE. By analogy we would expect antibodies to the RBD that block attachment to ACE2 to behave in a similar way to antibodies against the globular head of HA, whilst antibodies such as CR3022, that neutralise by an alternative mechanism to blocking receptor attachment, may need to be present throughout the incubation period with the indicator cells to reveal neutralisation. This agrees with our observation that, in the absence of ACE2, the CR3022 Fab destroys the prefusion-stabilised trimer (T_1/2_ ∼1h at room temperature as measured by cryo-EM).

With monoclonal antibodies now recognised as potential antivirals (Lu et al., 2020b; Salazar et al., 2017) our results suggest that CR3022 may be of immediate utility, since the mechanism of neutralisation will be unusually resistant to virus escape. In contrast antibodies which compete with ACE2 (whose epitope on SARS-CoV-2 is reported to have already shown mutation at residue 495 (GISAID: Accession ID: EPI_ISL_429783 Wienecke-Baldacchino et al., 2020 (Shu and McCauley, 2017)), are likely to be susceptible to escape. Furthermore, with knowledge of the detailed structure of the epitope presented here a higher affinity version of CR3022 might be engineered. Alternatively, since the same mechanism of neutralisation is likely to be used by other antibodies, a more potent monoclonal antibody targeting the same epitope might be found (for instance by screening for competition with CR3022). Additionally, since this epitope is sterically and functionally independent of the well-established receptor-blocking neutralising antibody epitope there is considerable scope for therapeutic synergy between antibodies targeting the two epitopes (indeed this type of synergy has been described for SARS-CoV (Ter Meulen et al., 2006)). Moreover, it has been reported (Wan et al., 2019) that antibody mediated enhancement occurs via antibodies that mimic receptor attachment whereas CR3022-like binding might circumvent this by pre-attachment conversion to the post-fusion state. Finally, display of this epitope on an RBD-based vaccine antigen might focus immune responses, conceivably mitigating the immunopathology reported for SARS-CoV (Perlman and Dandekar, 2005; Tseng et al., 2012).

## Method Details

### Cloning

#### CR3022

Two vectors were constructed containing resident human Cκ and IgG1 CH1 sequences and a signal sequence. Synthetic genes encoding the constant regions were inserted by Infusion® cloning into PmeI-HindIII cut pOPING-ET (Nettleship et al., 2008). The vectors have been engineered so that VL and VH sequences can be inserted into the KpnI-BsiWI (pOPINhuVL) and KpnI-SfoI (pOPINhuVH) restriction sites by Infusion® cloning. Synthetic genes encoding the candidate variable regions of CR3022 (Ter Meulen et al., 2006) were purchased from IDT Technologies (Leuven, Belgium) as gBlocks. The VH gene was amplified usingthe forward primer:5’-GGTTGCGTAGCTGGTACCCAGATGCAGCTGGTGCAATC-3’ and the reverse primer: 5’-GCCCTTGGTGGAGGCGACGGTGACCGTGGTCCCTTG; the VL gene was amplified using the forward primer 5’-GGTTGCGTAGCTGGTACCGACATCCAGTTGACCCAGTC-3’ and the reverse primer 5’-GTGCAGCCACCGTACGTTTGATTTCCACCTTGGTCCC-3’. The genes were inserted into the pOPIN expression vectors by Infusion® cloning.

The CR3022 hIgG1 heavy chain gene was amplified through joining three fragments (using the forward primer 5’-GCGTAGCTGAAACCGGCCAGATGCAGCTGGTGCAATC-3’ and the reverse primer 5’-GCCCTTGGTGGAGGCGCTAGAGACGGTGACCGTGGTCCCTTG-3’, and the CR3022 VH as template; the forward primer 5’-CAAGGGACCACGGTCACCGTCTCTAGCGCCTCCACCAAGGGC-3’ and the reverse primer 5’-CGGTGGGCATGTGTGAGTTTTGTCACAAGATTTGGGCTCAAC-3’, and the CR3022 VH as template; the forward primer 5’-GTTGAGCCCAAATCTTGTGACAAAACTCACACATGCCCACCG-3’ and the reverse primer 5’-GTGATGGTGATGTTTACCCGGAGACAGGGAGAGGCTCTTCTG-3’, and the pOPINTTGneoFc as template) using the forward primer 5’-GCGTAGCTGAAACCGGCCAGATGCAGCTGGTGCAATC-3’ and the reverse primer 5’-GTGATGGTGATGTTTACCCGGAGACAGGGAGAGGCTCTTCTG-3’. The gene was inserted into the vector pOPINTTGneo (Nettleship et al., 2015) incorporating a C-terminal His6 tag.

#### CR3022 used for neutralisation

The heavy and kappa light variable genes of the antibody were sourced from the Genbank ABA54613.1 and ABA54614.1 respectively and the codon optimised sequences were synthesized by GeneArt. These sequences were cloned into Antibody expression vectors (Genbank FJ475055 and FJ475056). Antibody was expressed using ExpiCHO expression system (LifeTechnologies) according to the manufacturer’s protocol and purified using a Protein A MabSelect SuRE column (GE Healthcare). The wash buffer contained 20mM Tris & 150mM NaCl buffered to pH 8.6 and the elution was done using 0.1 M Citric acid pH 2.5. The eluate was neutralised immediately using 1.5 M Tris pH 8.6 and then buffer exchanged to PBS using a 15 ml 30 kDa MWCO centrifugal filter (Merck Millipore).

#### RBD

The gene encoding amino acids 330-532 of the Receptor Binding Domain (RBD) of SARS-CoV-2 (Gene ID: MN908947) was amplified from a synthetic gene (IDT Technologies) using the forward primer 5’-GCGTAGCTGAAACCGGCCCGAATATCACAAATCTTTGTCC-3’ and the reverse primer 5’-GTGATGGTGATGTTTATTTGTACTTTTTTTCGGTCCGC-3’ or the reverse primer 5’-GTGATGGTGATGTTTTTCATGCCATTCAATCTTTTGTGCCTCAAAAATATCATTCAAATTTGTACTTTTTTTCGGTCCGC-3’ and inserted into the vector pOPINTTGneo incorporating either a C-terminal His6 or BirA-His6 tag.

#### ACE2

The gene encoding amino acids 19-615 of the human ACE2 was amplified from a an image clone (Sourcebiosciences, clone ID: 5297380) using the forward primer 5’-GCGTAGCTGAAACCGGCTCCACCATTGAGGAACAGGCC-3’ and the reverse primer 5’-GTGATGGTGATGTTTGTCTGCATATGGACTCCAGTC-3’ and inserted into the vector the vector pOPINTTGneo incorporating a C-terminal His6. The gene was also amplified using the forward primer 5’-GCGTAGCTGAAACCGGCTCCACCATTGAGGAACAGGCC-3’ and the reverse primer 5’-CAGAACTTCCAGTTTGTCTGCATATGGACTCCAGTC-3’ and inserted into the vector pOPINTTGneoFc incorporating a C-terminal hIgG1Fc-His6 tag.

#### Spike ectodomain

The gene encoding amino acids 1-1208 of the SARS-CoV-2 spike glycoprotein ectodomain, with mutations of RRAR > GSAS at residues 682-685 (the furin cleavage site) and KV > PP at residues 986-987, as well as inclusion of a T4 fibritin trimerisation domain, a HRV 3C cleavage site, a His-6 tag and a Twin-Strep-tag at the C-terminus. As reported by Wrapp *et al*. (Wrapp et al., 2020)

#### Validation and protein production

All vectors were sequenced to confirm clones were correct. Recombinant RBD, ACE2, CR3022 Fab and CR3022 IgG were transiently expressed in Expi293™ (ThermoFisher Scientific, UK) and proteins were purified from culture supernatants by an immobilised metal affinity using an automated protocol implemented on an ÄKTAxpress (GE Healthcare, UK) (Nettleship et al., 2009), followed by a Hiload 16/60 superdex 75 or a Superdex 200 10/300GL column, using phosphate-buffered saline (PBS) pH 7.4 buffer. Recombinant Spike ectodomain was expressed by transient transfection in HEK293S GnTI-cells (ATCC CRL-3022) for 9 days at 30 °C. Conditioned media was dialysed against 2x phosphate buffered saline pH 7.4 buffer. The Spike ectodomain was purified by immobilised metal affinity chromatography using Talon resin (Takara Bio) charged with cobalt followed by size exclusion chromatography using HiLoad 16/60 Superdex 200 column in 150 mM NaCl, 10 mM HEPES pH 8.0, 0.02% NaN_3_ at 4 °C, before buffer exchange into 2 mM Tris pH 8.0, 200 mM NaCl (Wrapp et al., 2020).

### Surface plasmon resonance

Surface plasmon resonance experiments were performed using a Biacore T200 (GE Healthcare). All assays were performed with a running buffer of PBS pH 7.4 supplemented with 0.005% v/v Surfactant P20 (GE Healthcare) at 25 °C. To determine the binding kinetics between the RBD of SARS-CoV-2 and CR3022 mAb, two different experimental settings were attempted. The first experiment was performed with the use of a CAP sensor chip (GE Healthcare). Biotin CAPture Reagent provided in the Biotin CAPture Kit (GE Healthcare) was captured onto the sensor chip according to manufacturer’s instructions. The RBD with a BirA tag was biotinylated using a biotinylation kit (Avidity, LLC) and was immobilized through the Biotin CAPture Reagent, at a density of 15-30 RU on the sample flow cell. The reference flow cell was left blank. The CR3022 Fab was injected over the two flow cells at a range of five concentrations prepared by serial two-fold dilution from 95 nM, at a flow rate of 30 μL/min using a Single-cycle kinetics program with an association time of 60 s and a dissociation time of 60 s. Running buffer was also injected using the same program for background subtraction. The second experiment was performed using a Sensor Chip Protein A (GE Healthcare). CR3022 IgG was immobilised at a density of approximately 30 RU on the sample flow cell. The reference flow cell was left blank. The RBD was injected over the two flow cells at a range of five concentrations prepared by serial two-fold dilution from 100 nM, at a flow rate of 30 μL/min using a Single-cycle kinetics program with an association time of 75 s and a dissociation time of 60 s. Running buffer was also injected using the same program for background subtraction. All data were fitted to a 1:1 binding model using the Biacore T200 Evaluation Software 3.1. In the competition assay where CR3022 IgG was used as the ligand, approximately 1000 RU of CR3022 IgG was immobilised onto a Sensor Chip Protein A. The following samples were injected: (1) 1 µM ACE2, (2) 1 µM (anti-Caspr2) E08R Fab; (3) a mixture of 1 µM ACE2 and 0.1 µM RBD, (4) a mixture of 1 µM E08R Fab and 0.1 µM RBD, and (4) 0.1 µM RBD. In the competition assay where ACE2-hIgG1Fc was used as the ligand, approximately 1000 RU of ACE2-hIgG1Fc was immobilised onto a Sensor Chip Protein A. The following samples were injected: (1) 1 µM CR3022 Fab, (2) 1 µM E08R Fab; (3) a mixture of 1 µM CR3022 Fab and 0.1 µM RBD, (4) a mixture of 1 µM E08R Fab and 0.1 µM RBD, and (4) 0.1 µM RBD. All injections were performed with an association time of 60 s and a dissociation time of 600 s. All curves were plotted using GraphPad Prism 8.

### Bio-layer interferometry

To further validate the SPR results the K_D_ of Fab CR3022 for RBD was also measured by bio-layer interferometry. Kinetic assays were performed on an Octet Red 96e (ForteBio) at 30 °C with a shake speed of 1000 rpm. Fab CR3022 was immobilized onto amine reactive 2nd generation (AR2G) biosensors (ForteBio) and serially diluted RBD (80,40,20,10 and 5 nM) was used as analyte. PBS (pH 7.4) was used as the assay buffer. Recorded data were analysed using the Data Analysis Software HT v11.1 (Fortebio), with a global 1:1 fitting model.

### Neutralisation

Neutralising virus titres were measured in serum samples that had been heat-inactivated at 56 °C for 30 minutes. SARS-CoV-2 (<u>strain Victoria/1/2020 at cell passage 3</u>(Caly et al., 2020)) was diluted to a concentration of 1.4E+03 pfu/mL (70 pfu/50 µl) and mixed 50:50 in 1% FCS/MEM containing 25 mM HEPES buffer with doubling serum dilutions from 1:10 to 1:320 in a 96-well V-bottomed plate.

The plate was incubated at 37 °C in a humidified box for 1 hour to allow the antibody in the serum samples to neutralise the virus. CR3022 (pH7.2) at a starting concentration of 2 mg/mL was diluted 1 in 10. The dilutions were then made 2-fold up to 320. The neutralised virus was transferred into the wells of a twice DPBS-washed plaque assay 24-well plate that had been seeded with Vero/hSLAM the previous day at 1.5E+05 cells per well in 10% FCS/MEM. Neutralised virus was allowed to adsorb at 37 °C for a further hour, and overlaid with plaque assay overlay media (1X MEM/1.5% CMC/4% FCS final). After 5 days incubation at 37 °C in a humified box, the plates were fixed, stained and plaques counted. Dilutions and controls were performed in duplicate. Median neutralising titres (ND50) were determined using the Spearman-Karber formula (Kärber, 1931) relative to virus only control wells.

### Crystallization, data collection and X-ray structure determination

Purified and deglycosylated RBD and CR3022 Fab were concentrated to 8.3 mg/mL and 11 mg/mL respectively, and then mixed in an approximate molar ratio of 1:1. Crystallization screen experiments were carried out using the nanolitre sitting-drop vapour diffusion method in 96-well plates as previously described (Walter et al., 2003, 2005). Crystals were initially obtained from Hampton Research PEGRx HT screen, condition 63 containing 0.1 M Sodium malonate, 0.1 M Tris pH 8.0 and 30% w/v Polyethylene glycol 1,000. The best crystals were grown in drops containing 200 nl sample and 100 nl reservoir solution.

Crystals were mounted in loops and frozen in liquid nitrogen prior to data collection. Diffraction data were collected at 100 K at beamline I03 of Diamond Light Source, UK. Diffraction images of 0.1° rotation were recorded on an Eiger2 XE 16M detector (exposure time of either 0.002 s or 0.01 s per frame, beam size 80×20 μm and 100% beam transmission). Data were indexed, integrated and scaled with the automated data processing program Xia2-dials (Winter, 2010; Winter et al., 2018). The data set of 720° was collected from a single frozen crystal to 4.4 Å resolution with 52-fold redundancy. The crystal belongs to space group *P4*_*1*_*2*_*1*_*2* with unit cell dimensions *a* = *b* = 150.5 Å and *c* = 241.6 Å. The structure was determined by molecular replacement with PHASER (McCoy et al., 2007) using search models of human germline antibody Fabs 5-51/O12 (PDB ID, 4KMT (Teplyakov et al., 2014)) heavy chain and IGHV3-23/IGK4-1 (PDB ID, 5I1D (Teplyakov et al., 2016)) light chain, and RBD of SARS-CoV-2 RBD/ACE2 complex (PDB ID, 6M0J (Lan et al., 2020)). There is one RBD/CR3022 complex in the crystal asymmetric unit, resulting in a crystal solvent content of ∼87%.

During optimization of the crystallization conditions, a second crystal form was found to grow in the same condition with similar morphology. A data set of 720° rotation with data extending to 2.4 Å was collected on beamline I03 of Diamond from one of these crystals (exposure time 0.004 s per 0.1° frame, beam size 80×20 μm and 100% beam transmission). The crystal also belongs to space group *P4*_*1*_*2*_*1*_*2* but with significantly different unit cell dimensions (*a* = *b* = 163.1 Å and *c* = 189.1 Å). There were two RBD/CR3022 complexes in the asymmetric unit and a solvent content of ∼74%.

### X-ray crystallographic refinement and electron density map generation

The initial structure was determined using the lower resolution data from the first crystal form. Data were excluded at a resolution below 35 Å as these fell under the beamstop shadow. One cycle of REFMAC5 (Murshudov et al., 2011) was used to refine atomic coordinates after manual correction in COOT (Emsley and Cowtan, 2004) to the protein sequence from the search model. The software suite Vagabond was used to convert the atomic model into a bond-based description suited for low resolution refinement (Ginn, submitted). This described the protein model through a series of identical but positionally displaced conformers (referred to as an ensemble). The flexibility was described through whole-molecule translations and rotations per polypeptide chain and intramolecular flexibility through variation in torsion angles of bonds connecting C-alpha atoms. These torsion variations were constrained, with bonds of a similar effect on the flexibility of the protein structure moving in tandem. A global B factor of 130 was applied to the model to account for most of the disorder in the crystal. Alternate rounds of refinement were performed of (a) these flexibility parameters and (b) rigid body refinement of each polypeptide chain, for both the target function was the correlation coefficient with the electron density in real space. Local adjustments of atoms were performed in COOT (Emsley and Cowtan, 2004) using the Vagabond map and average model output coordinates. After local real-space refinement, updated coordinates were reloaded into Vagabond and bond torsion angles were adjusted to match them. Best electron density maps accounting for sources of phase error were output as a list of Fourier coefficients. Maps were sharpened by applying a B factor of −100 (Figure S5). The final refined structure had an R_work_ of 0.331 (R_free_, 0.315) for all data to 4.36 Å resolution. This structure was later used to determine the structure of the second crystal form, which has been refined with PHENIX (Liebschner et al., 2019) to R_work_ = 0.213 and R_free_ = 0.239 for all data to 2.42 Å resolution. This refined model revealed the presence of one extra residue at each heavy chain N-terminus and 3 extra residues at the N-terminus of one RBD from the signal peptide. There is well ordered density for a single glycan at each of the glycosylation sites at N331 and N343 in one RBD, and only one at N343 in the second RBD.

Data collection and structure refinement statistics are given in Table S3. Structural comparisons used SHP (Stuart et al., 1979), residues forming the RBD/Fab interface were identified with PISA (Krissinel and Henrick, 2007), figures were prepared with PyMOL (The PyMOL Molecular Graphics System, Version 1.2r3pre, Schrödinger, LLC).

### CR3022 Fab complex preparation and cryo-EM data collection

Purified spike protein was buffer exchanged into 2 mM Tris pH 8.0, 200 mM NaCl, 0.02 % NaN3 buffer using a desalting column (Zeba, Thermo Fisher). A final concentration of 0.2 mg/mL was incubated with CR3022 Fab (in the same buffer) in a 6:1 molar ratio (Fab to trimeric spike) at room temperature. Aliquots were taken at 50 minutes and 3 h. Immediately an aliquot was taken 3 μL of it was applied to a holey carbon-coated 200mesh copper grid (C-Flat, CF-2/1, Protochips) that had been freshly glow discharged on high for 20 s (Plasma Cleaner PDC-002-CE, Harrick Plasma) and excess liquid removed by blotting for 6 s with a blotting force of −1 using vitrobot filter paper (grade 595, Ted Pella Inc.) at 4.5 °C, 100 % relative humidity. Blotted grids were then immediately plunge frozen using a Vitrobot Mark IV (Thermo Fisher).

Frozen grids were first screened on a Glacios microscope operating at 200 kV (Thermo Fisher) before imaging on a Titan Krios G2 (Thermo Fischer) at 300 kV. Movies (40 frames each) were collected in compressed tiff format on a K3 detector (Gatan) in super resolution counting mode using a custom EPU version 2.5 (Thermo Fischer) with a defocus range of 0.8-2.6 μm and at a nominal magnification of x105,000, corresponding to a calibrated pixel size of 0.83 Å/pixel, see Table S4.

### Cryo-EM data processing

For both the 50 minute and 3 h incubation datasets, motion correction and alignment of 2x binned super-resolution movies was performed using Relion3.1. CTF-estimation with GCTF (v1.06) (Zhang, 2016) and non-template-driven particle picking was then performed within cryoSPARC v2.14.1-live followed by multiple rounds of 2D classification (Punjani et al., 2017).

For the 50 minutes dataset. 2D class averages for structure-A and structure-B were then used separately for template-driven classification before further rounds of 2D and 3D classification with C1 symmetry. Both structures were then sharpened in cryoSPARC. Data processing and refinement statistics are given in Table S4.

An initial model for the spike (structure-A) was generated using PDB ID, 6VYB (Walls et al., 2020) and rigid body fitted into the final map using COOT (Emsley and Cowtan, 2004). The model was further refined in real space with PHENIX (Liebschner et al., 2019) which resulted in a correlation coefficient of 0.84. Two copies of RBD-CR3022 were fitted into structure-B in the same manner. Because of the strongly anisotropic resolution the overall correlation coefficient vs the model was lower (0.47).

For the 3 h incubation dataset, particles were extracted with a larger box size (686 pixels as compared to 540 pixels), and, following multiple rounds of 2D classification, 2D class averages from ‘blob-picked’ particles showing signs of complete ‘flower-like’ structures were selected for *ab initio* reconstruction. For the 3 h data no detailed fitting was attempted.

## Acknowledgements

This work was supported by a grant from the CAMS-Oxford Institute to D.I.S. E.E.F and J.Ren are supported by the Wellcome Trust (101122/Z/13/Z), Y.Z. by Cancer Research UK (C375/A17721) and D.I.S. and E.E.F. by the UK Medical Research Council (MR/N00065X/1). J.H. is supported by a grant from the EPA Cephalosporin Fund. PPUK is funded by the Rosalind Franklin Institute EPSRC Grant no. EP/S025243/1. The National Institute for Health Research Biomedical Research Centre Funding Scheme supports G.R.S. together with the Chinese Academy of Medical Sciences (CAMS) Innovation Fund for Medical Science (CIFMS), China (grant number: 2018-I2M-2-002), which also supports D.I.S. G.R.S. is also supported as a Wellcome Trust Senior Investigator (grant 095541/A/11/Z). T.M. is supported by Cancer Research UK grants C20724/A14414 and C20724/A26752 to Christian Siebold. This is a contribution from the UK Instruct-ERIC Centre. The Wellcome Centre for Human Genetics is supported by the Wellcome Trust (grant 090532/Z/09/Z). Virus used for the neutralisation assays was a gift from Julian Druce, Doherty Centre, Melbourne, Australia. We acknowledge Diamond Light Source for time on Beamline I03 under Proposal mx19946 and for electron microscope time at the UK national electron bio-imaging centre (eBIC), Proposal BI26983-2, both COVID-19 Rapid Access. Huge thanks to the teams, especially at the Diamond Light Source and Department of Structural Biology, Oxford University that have enabled work to continue during the pandemic.

## Author Information

These authors contributed equally: Y. Zhao, J. Huo.

## Contributions

J.H. and D.Z. performed interaction analyses and T.T., P.R., A.T., N.C., K.B. and M.C. prepared material for, and executed, neutralisation assays. Y.Z., J.H., J.Ren, performed sample preparation for and crystallographic experiments and processed the data. N.G.P. assisted with X-ray diffraction data collection. H.G. and J.Ren. refined the structures and together with E.E.F. and D.I.S. analysed the results. G.R.S., J.M., and P.S. prepared the Spike construct. L.C. helped performed cryo-EM data processing, T.M., R.R.R. and P.N.M.S. prepared the spike sample, H.M.E.D. performed cryo-EM sample preparation screening and processing and J.Radecke performed cryo-EM data collection. E.E.F., J.Ren, Y.Z. and D.I.S. wrote the manuscript. All authors read and approved the manuscript.

## Competing interests

The authors declare no competing interests.

## Corresponding authors

Correspondence to Jingshan Ren or David I. Stuart.

## Data availability

The high resolution and lower resolution coordinates and structure factors of the SARS-CoV-2 RBD/CR3022 complex are available from the PDB with accession codes 6YLA and 6YM0 respectively. EM maps and structure models are deposited in EMDB and PDB with accession codes XXX and YYY for the prefusion Spike, and EMD-10863 and 6YOR for the dimeric RBD/CR3022 complex respectively. The data that support the findings of this study are available from the corresponding authors on request.

## Notes

### Competing Interest Statement

The authors have declared no competing interest.

