## Supplemental Information for "Neutralization of SARS-CoV-2 by destruction of the prefusion Spike"

Figures S1-S10.

Tables S1-S4.

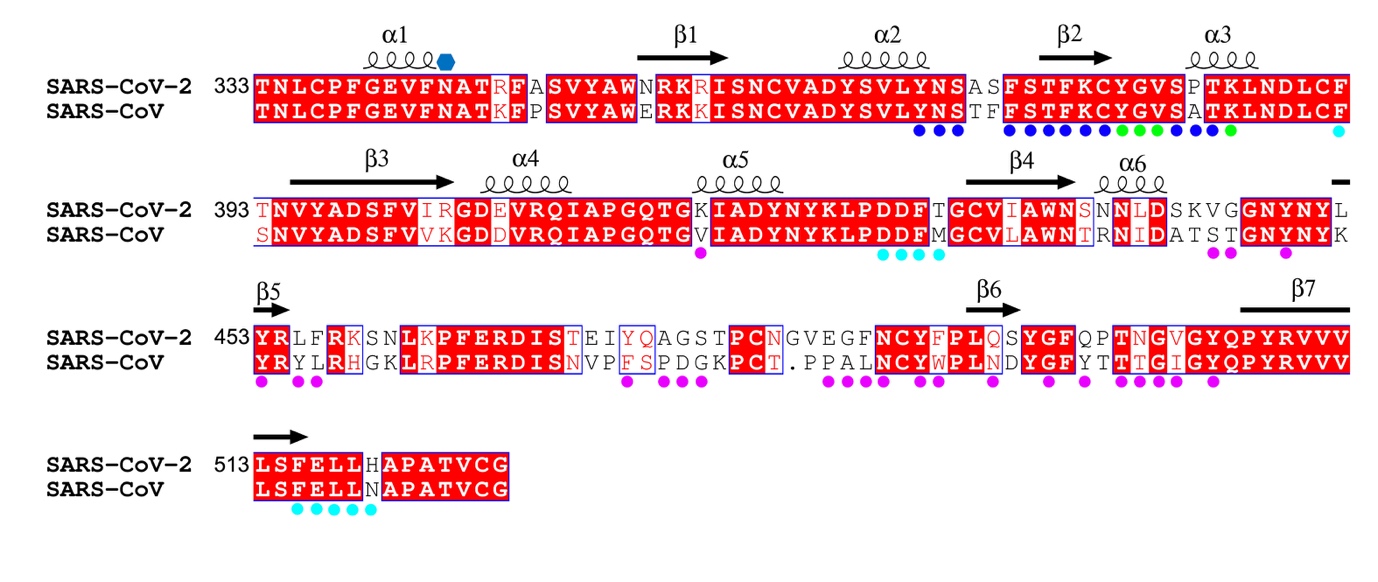

**Figure S1 | Sequence alignment between the RBDs of SARS-CoV and SARS-CoV-2.** Residue numbers are those of SARS-CoV-2 RBD, conserved amino acids have a red background, secondary structures are labelled on the top of the sequence, and the glycosylation site is marked with a blue hexagon. Residues involved in receptor binding are marked with magenta disks. Blue disks mark the residues involved in interactions with the CR3022 heavy chain, cyan disks mark the residues interacting with the CR3022 light chain and green disks with both chains.

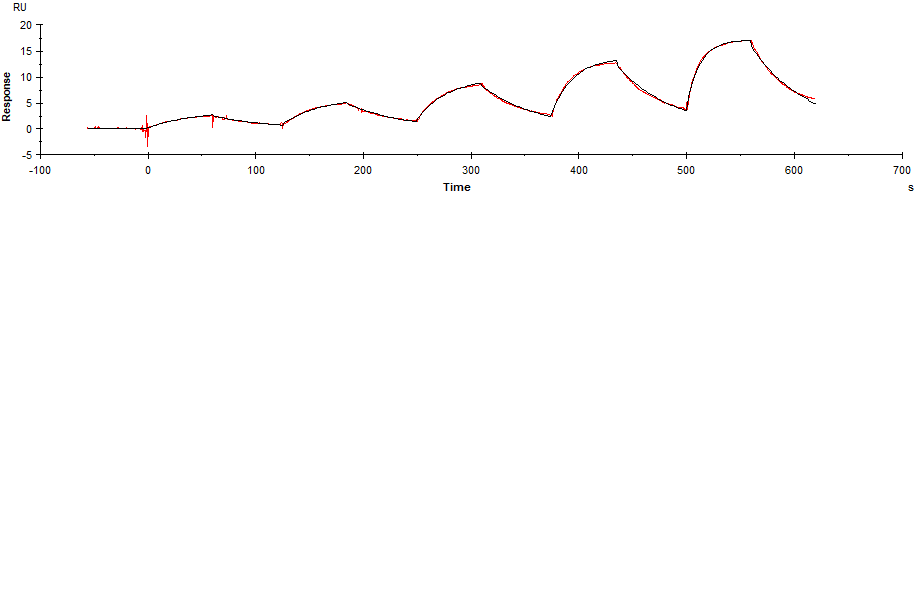
**a**

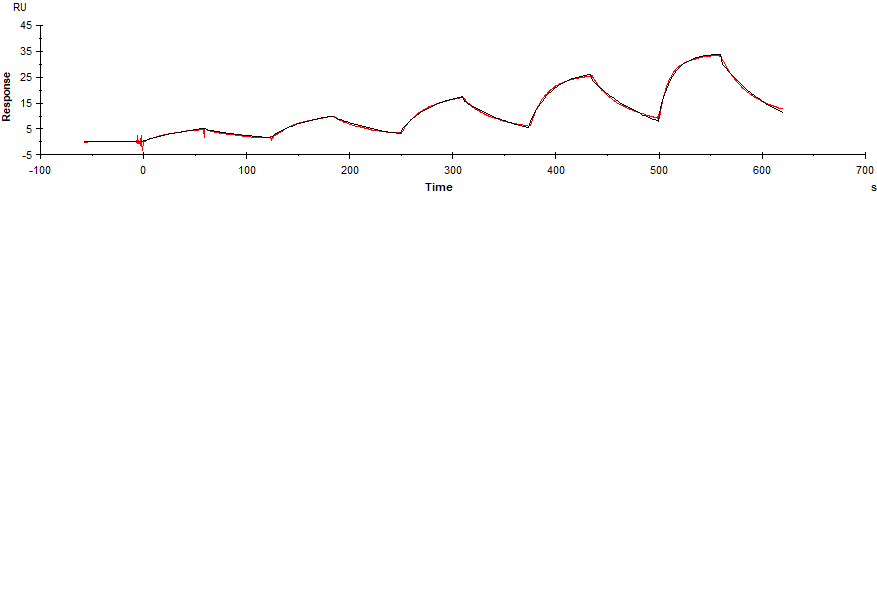
**b**

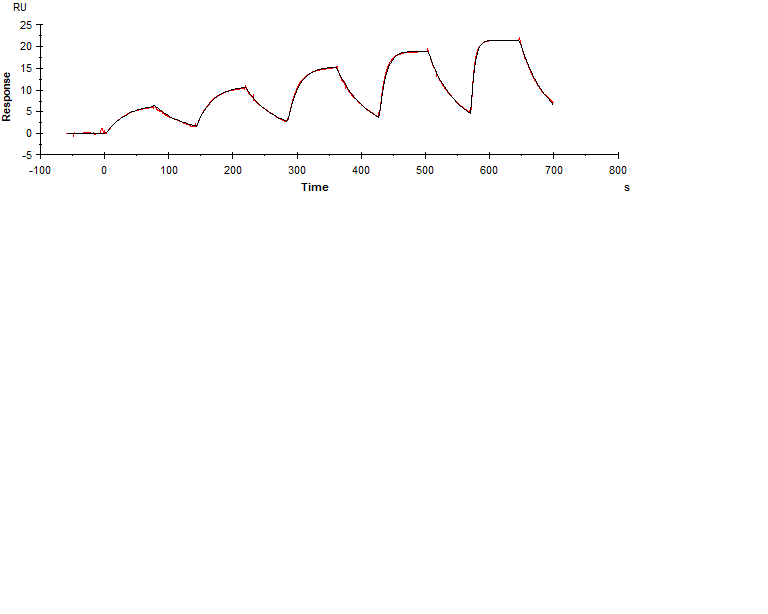
**c**

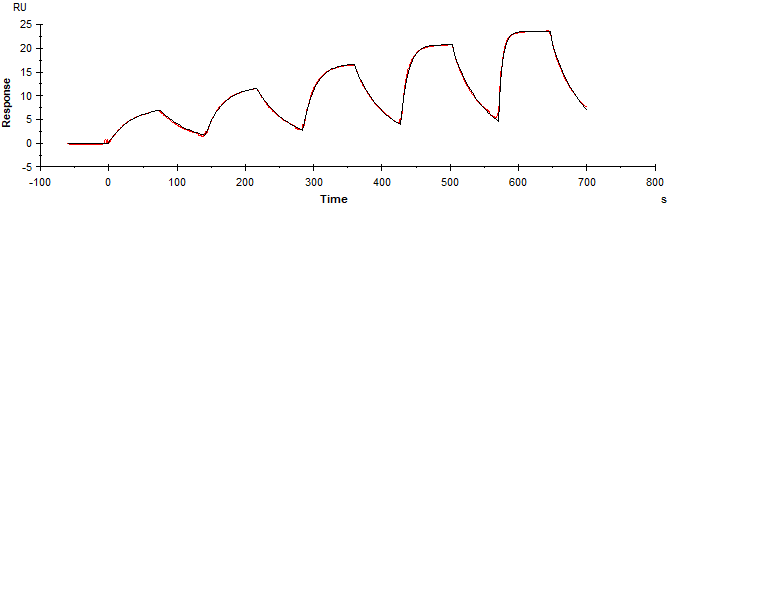
**d**

**e**

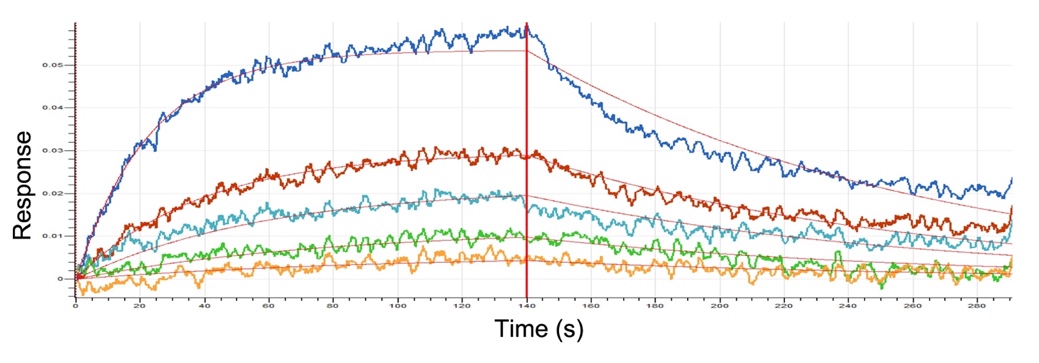

**Figure S2 | Binding affinity between RBD and CR3022 Fab. a-b,** Surface plasmon resonance binding sensorgrams measured with a Biacore T200. Biotinylated (Bio-) RBD was immobilised as the ligand and CR3022 Fab was used as analyte at five concentrations (5.9, 11.9, 23.8, 47.5 and 95 nM). **c-d**, CR3022 IgG was immobilised as the ligand and RBD-His was used as analyte at five concentrations (6.25, 12.5, 25, 50, 100 nM). Data were fitted to a 1:1 binding model using the Biacore T200 Evaluation Software 3.1. The average kinetic values from these two sets of experiment are listed in Extended Table 1. e**,** Binding sensorgram of the interaction between RBD and CR3022 Fab measured with an Octet platform. CR3022 Fab was immobilized onto AR2G biosensors, and RBD was used as analyte with a serial dilution of 5,10, 20, 40 and 80 nM. The measured K_D_ is 19 nM using a global 1:1 fitting model.

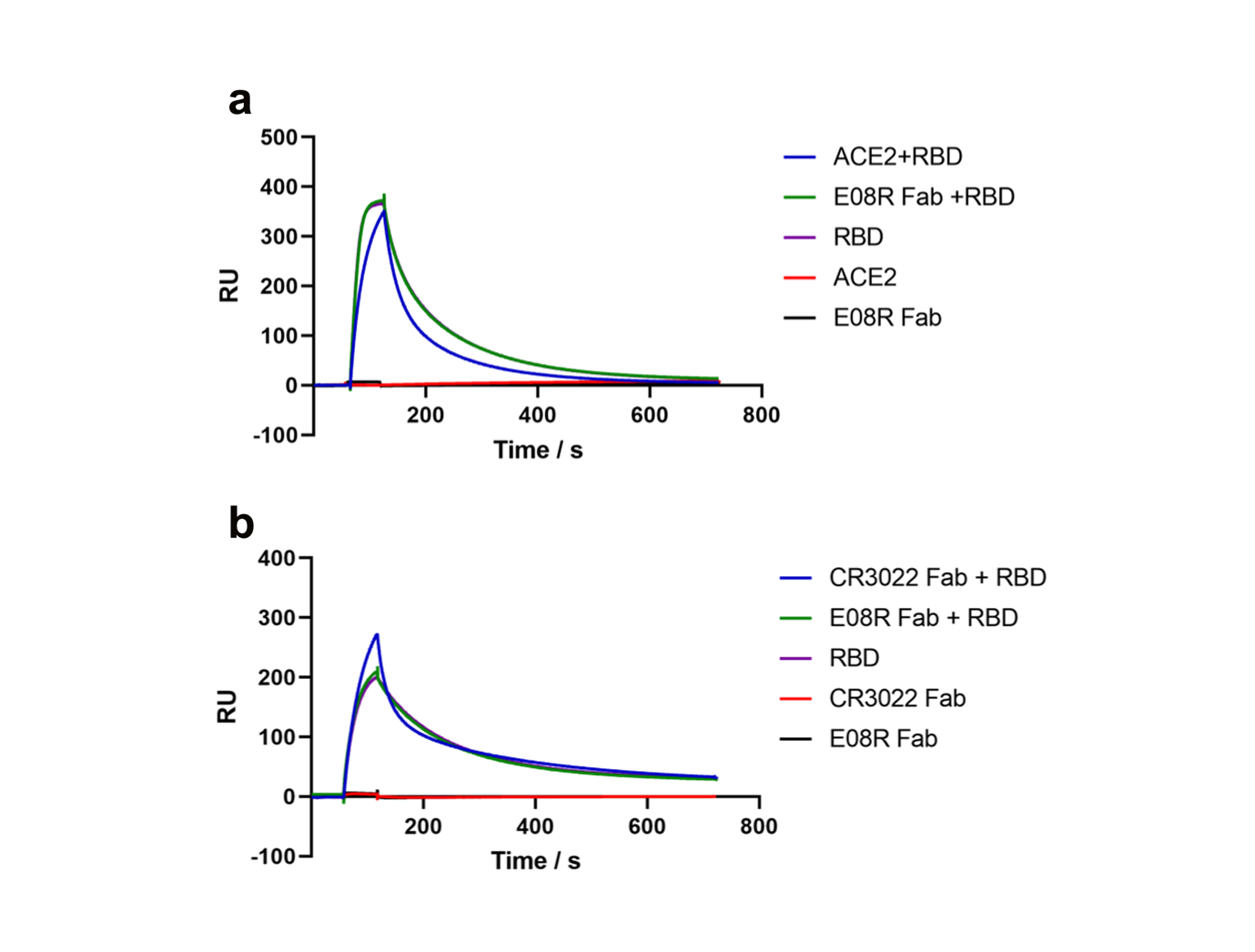

**Figure S3 | Binding competition of ACE2 and CR3022 Fab for RBD.** Surface plasmon resonance binding sensorgrams measured with a Biacore T200. **a,** CR3022 IgG was immobilised as the ligand, and the following samples were injected: (1) a mixture of 1 µM ACE2 and 0.1 µM RBD; (2) a mixture of 1 µM E08R Fab and 0.1 µM RBD; (3) ) 0.1 µM RBD; (4) 1 µM ACE2; (5) E08R Fab. **b,** ACE2-hIgG1Fc was immobilised as the ligand, and the following samples were injected: (1) a mixture of 1 µM CR3022 Fab and 0.1 µM RBD; (2) a mixture of 1 µM E08R Fab and 0.1 µM RBD; (3) 0.1 µM RBD; (4) 1 µM CR3022 Fab; (5) 1 µM E08R Fab.

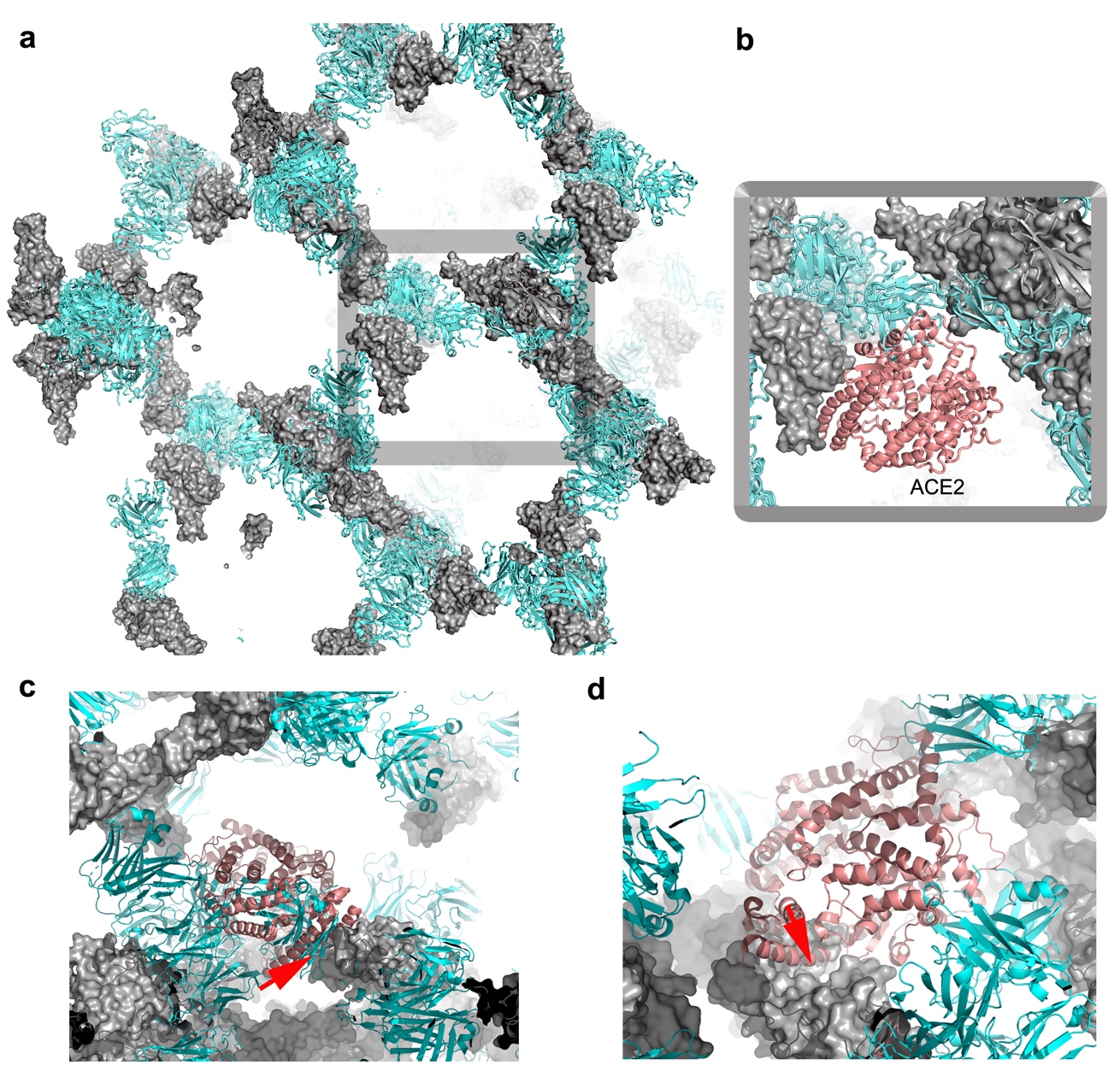

**Figure S4 | Crystal lattice of the RBD/CR3022 complex**. **a**, The packing of the RBD/CR3022 complex within the first crystal form. The RBD is shown as a grey surface and CR3022 Fab as cyan ribbons. **b**, A closeup of the crystal lattice with the RBD of the receptor complex overlapped onto the RBD of the Fab complex showing that the receptor binding site of the RBD is not blocked in the crystal. **c**, **d**, The ACE2 binding sites of the 2 RBDs in the second crystal form are blocked by crystal contact (indicated by red arrows).

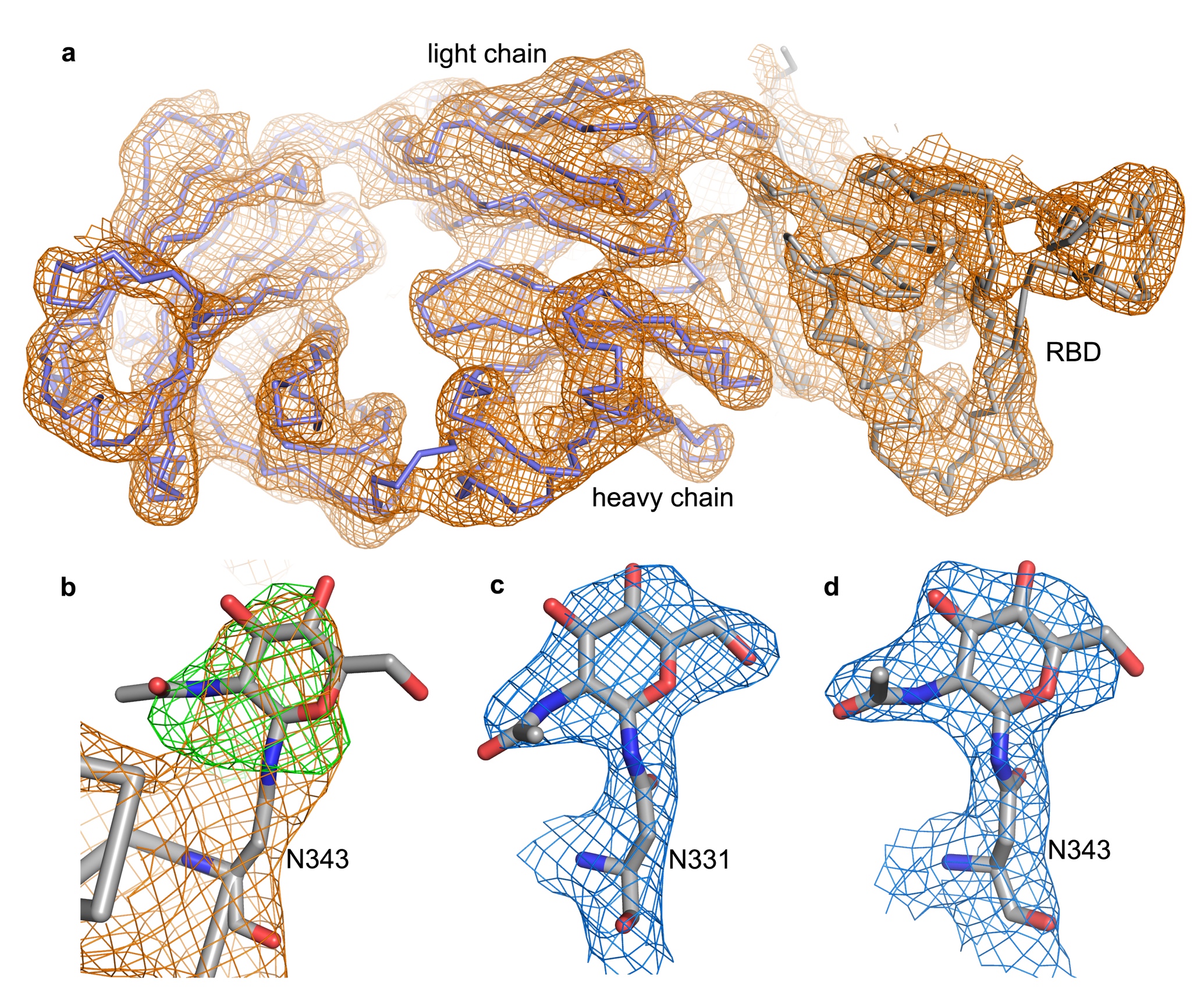

**Figure S5 | Electron density maps. a**, 4.4 Å resolution electron density map for crystal form 1, produced with Vagabond (ref) and contoured at 1.2 σ showing the overall quality of the structure. **b**, Difference electron density map (green) contoured at 3 σ showing the glycosylation site at N343 of the RBD. The glycan was not modelled into the structure used for the map calculation. **c**, **d**, Electron density maps of the glycosylation sites N331 (**c**) and N343 (**d**) in the second, high resolution (2.4 Å), crystal form.

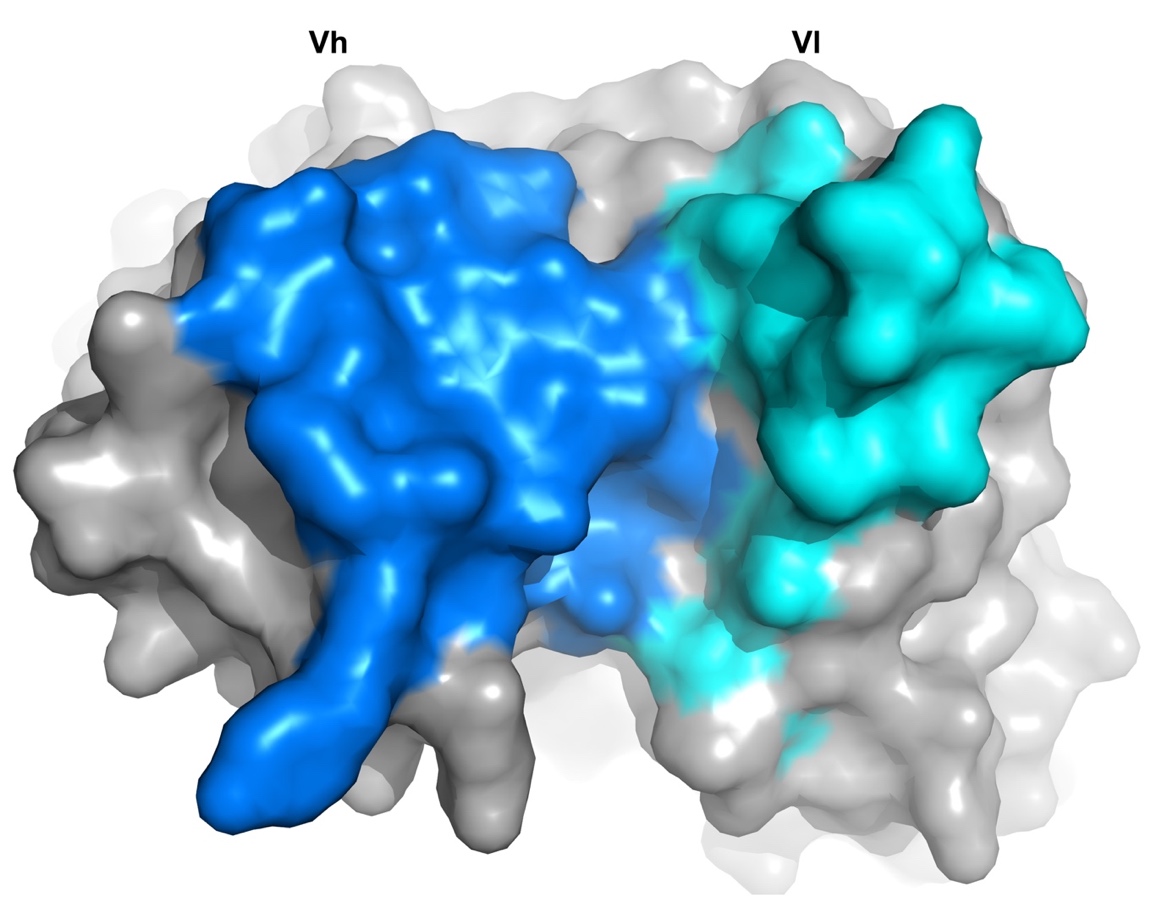

**Figure S6 | Buried solvent accessible area of the CR3022 antigen binding region due to engagement with the RBD.** Buried areas are coloured in blue for the heavy chain and cyan for the light chain.

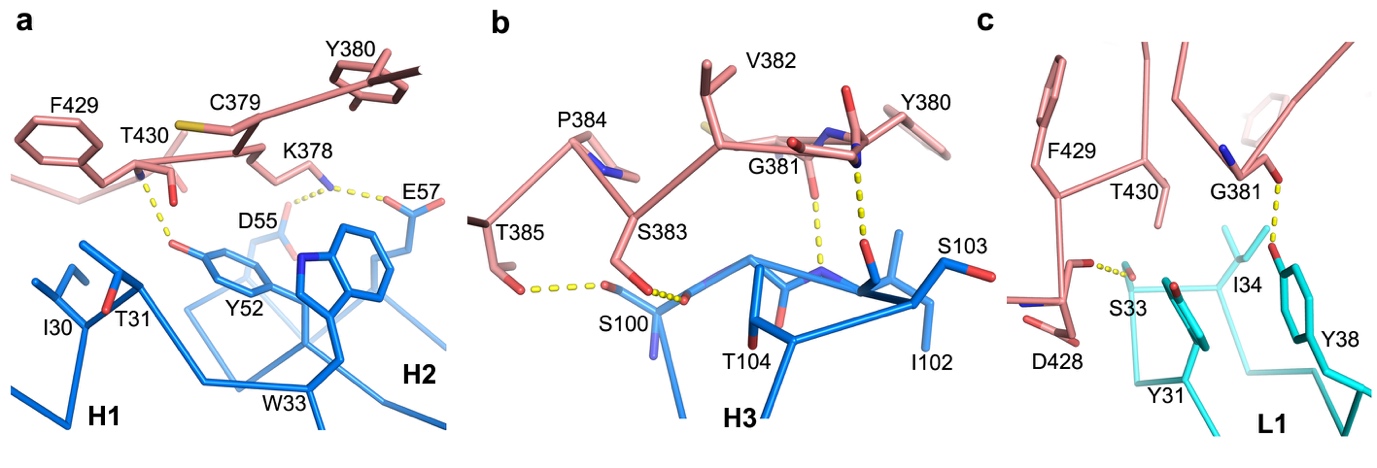

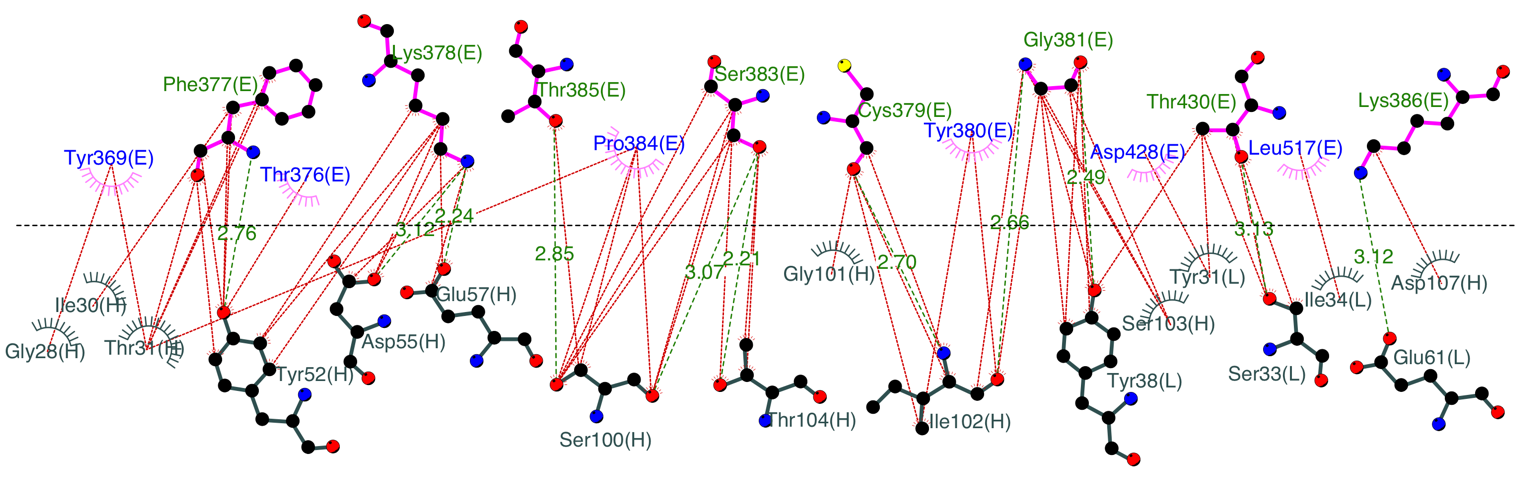
**d**

**Figure S7 | Details of contacts between the RBD and CR3022. a,** Contacts of the RBD with CR3022 heavy chain CDR1 (H1) and CDR2 (H2), and **b,** with CDR3 (H3). **c**, Interactions between the RBD and the light chain CDR1 (L1). Main chain backbones are shown as thinner sticks and side chains as thick sticks (RBD, salmon; heavy chain, blue; light chain, cyan). The yellow broken sticks represent hydrogen bonds or salt bridges. **d**, Ligplot (Laskowski and Swindells, 2011) representation of the interface details (chain identifiers: L, CR3022 light chain. H, CR3022 heavy chain, E, RBD.

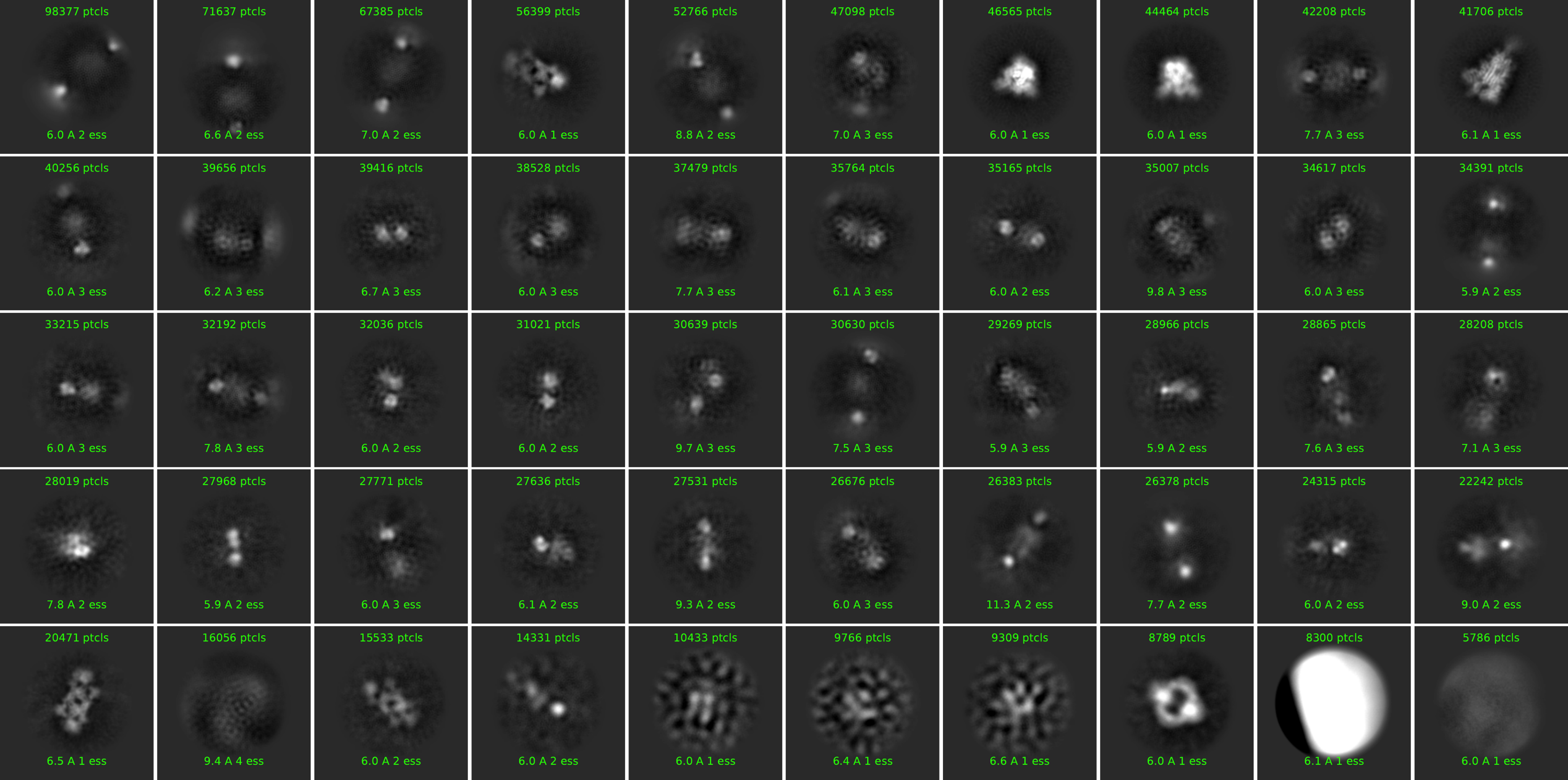
**a**

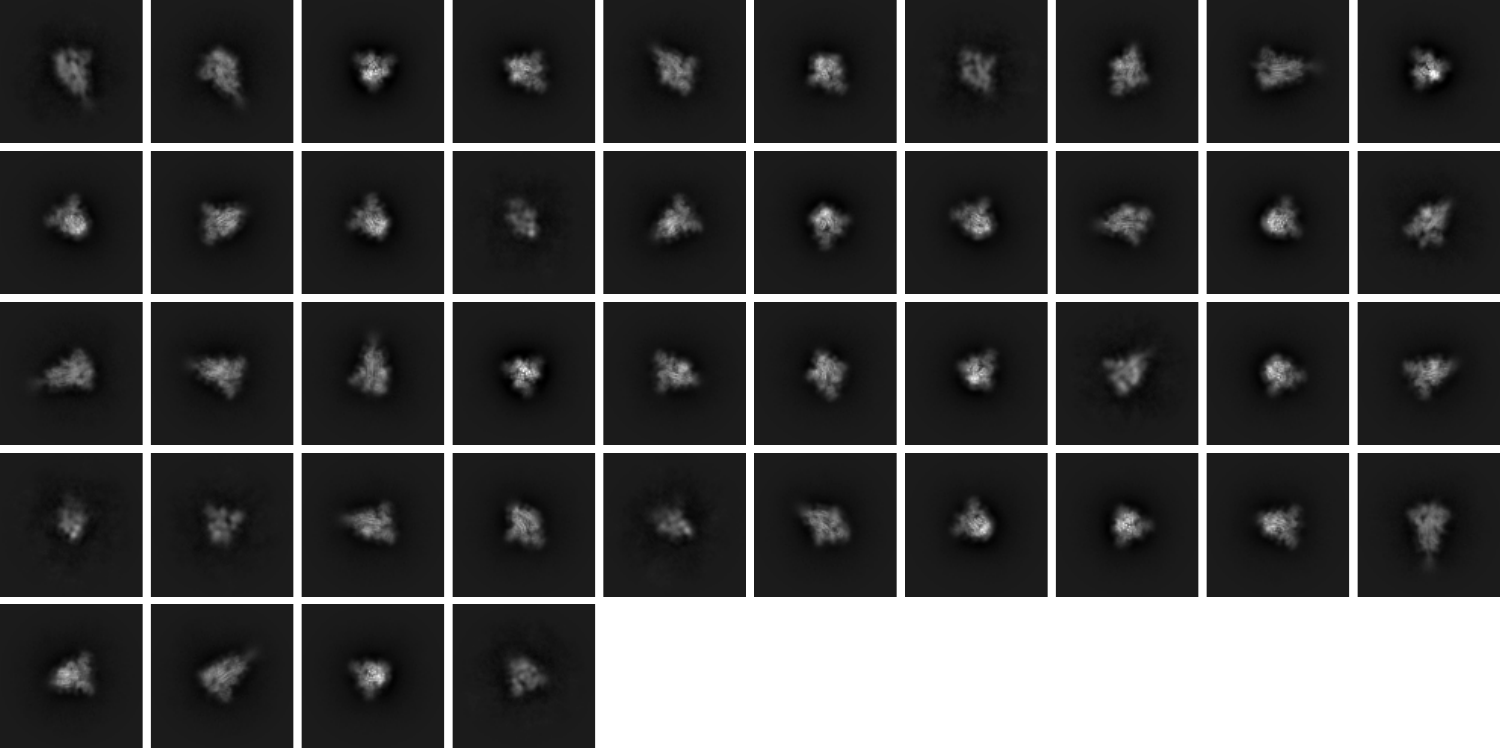
**b**

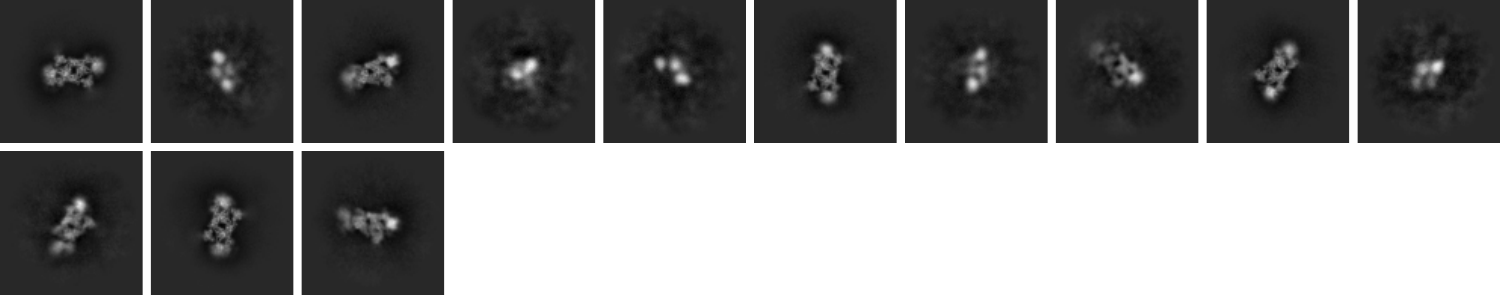
**c**

**Figure S8 | 2D class averages for 50 minutes incubation. a,** unbiased classes. **b**, selected classes for the prefusion state. **c**, selected classes for the dimeric association. See Methods for details.

**a**

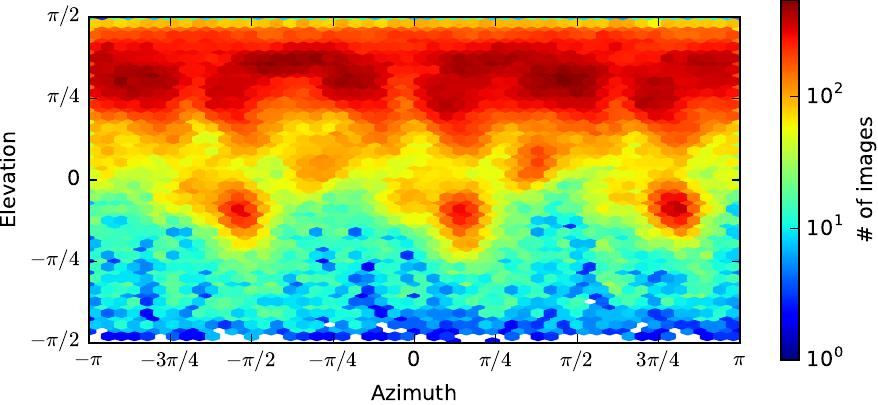

**b**

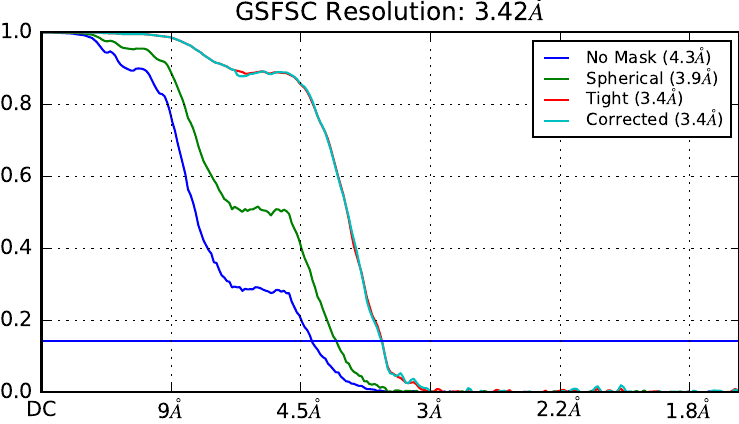

**Figure S9 | Prefusion Spike analysis for 50 minutes incubation. a,** orientation distribution. **b**, FSC analysis. See Methods for details.

**a**

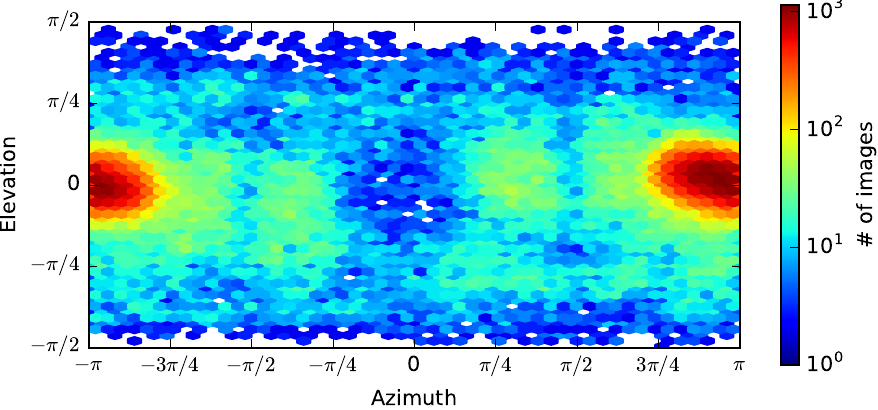

**b**

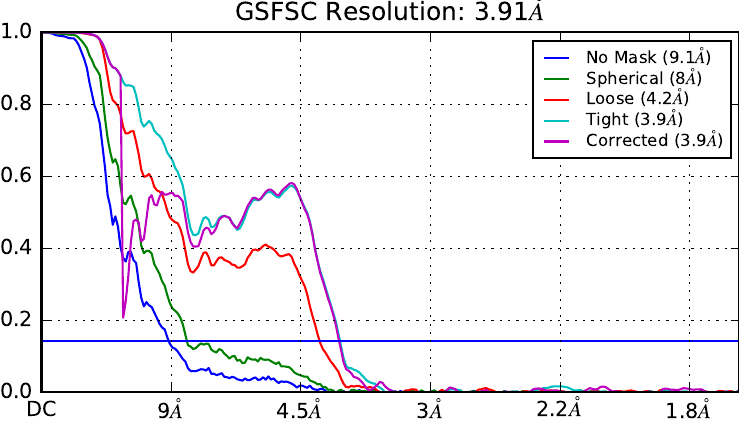

**Figure S10 | Dimeric CR3022/PBD analysis for 50 minutes incubation. a,** orientation distribution. **b**, FSC analysis. See Methods for details.

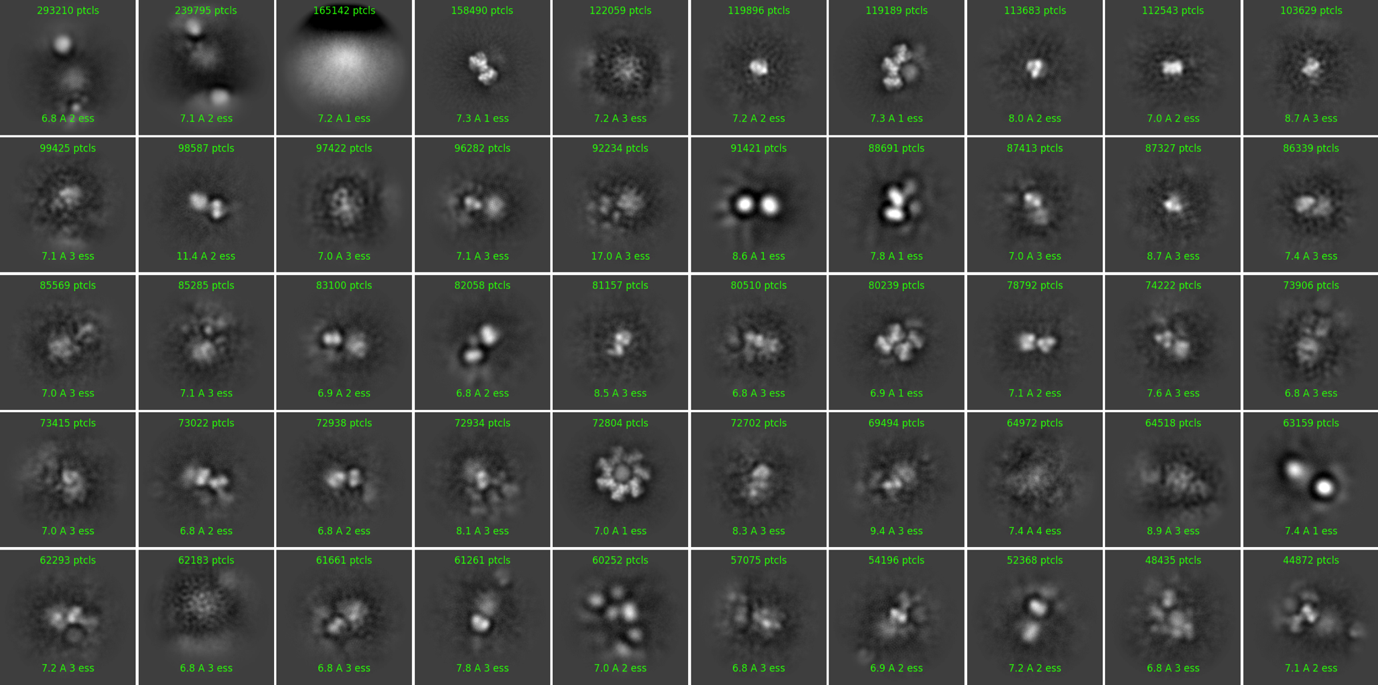
**a**

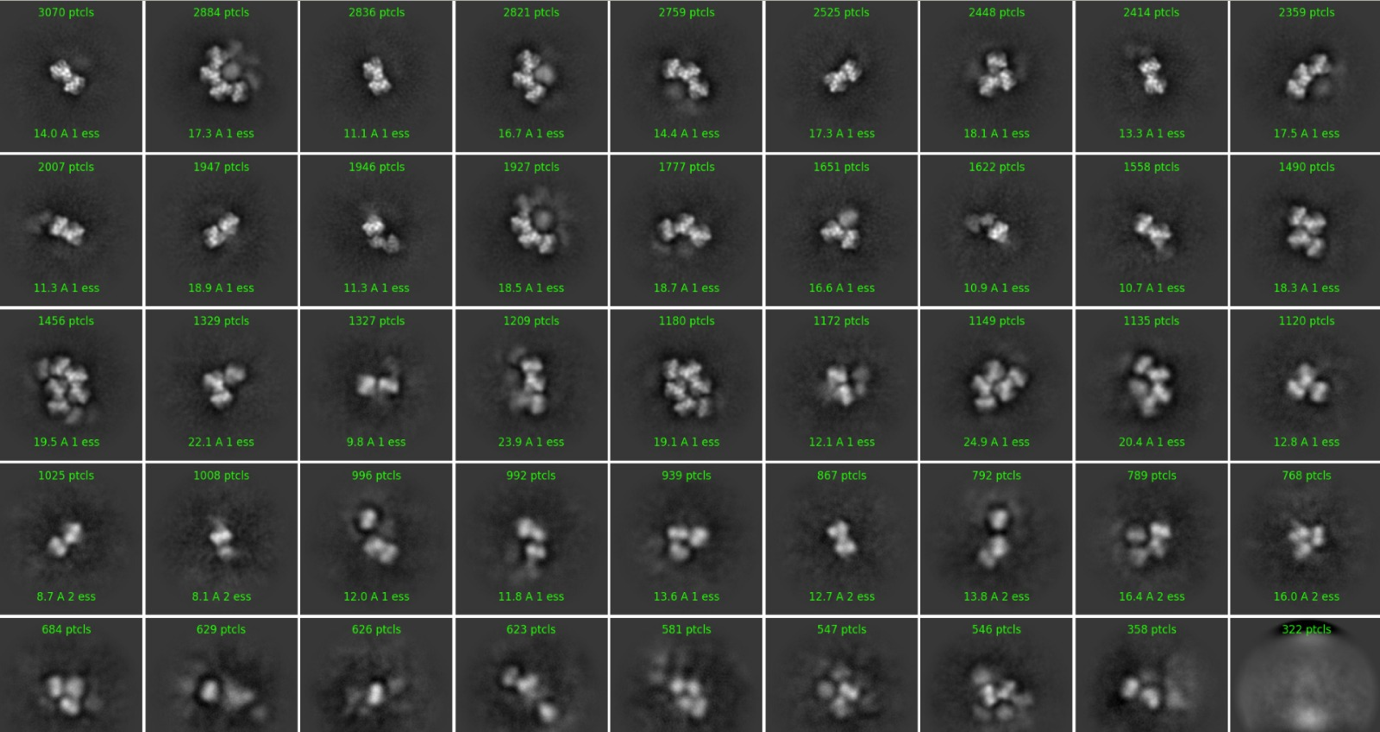
**b**

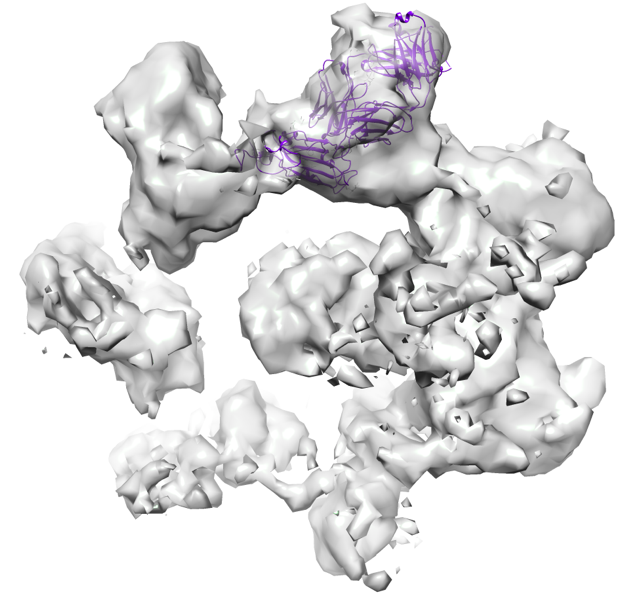
**c**

**Figure S11 | Analysis of cryo-EM data for 3 h incubation.** 2D class averages: **a,** unbiased classes. **b**, selected classes showing oligomeric assemblies. **c,** Ab initio model from 3 h incubation dataset to indicate how the CR3022 Fab/RBD complex might be accommodated within one oligomeric unit derived from classes in **b**.

**Table S1** | **SPR kinetic results**

| **Ligand** | **Biotinylated RBD** | **CR3022 IgG** |
| --- | --- | --- |
| Analyte | CR3022 Fab | His-tagged RBD |
| Ka (M^-1^s^-1^) | 6.3E+05 | 1.5E+06 |
| Kd (s^-1^) | 1.9E-02 | 2.3E-02 |
| KD (nM) | 30 | 15 |

**Table S2** | **Plaque Reduction Neutralization Test results**

| **ID** | **Description** | **ND50** |
| --- | --- | --- |
| Positive control | Convalescent serum | 1:149 |
| CR3022 | Mab | 1:201 |

**Table S3** | **X-ray data collection and refinement statistics**

| **Data collection** | | |
| --- | --- | --- |
| Data set | Crystal form 1 | Crystal form 2 |
| Space group | *P4_1_2_1_2* | *P4_1_2_1_2* |
| Cell dimensions (Å) | *a*=150.5, *b*=150.5, *c*=241.6 | *a*=163.1, *b*=163.1, *c*=189.1 |
| Resolution (Å) | 80.5–4.36 (4.44–4.36) | 58.8–2.42 (2.46–2.42) |
| Unique reflections | 18822 (931) | 97407 (4803) |
| *R*_merge_ | 0.683 (---) | 0.303 (---) |
| *R_pim_* | 0.097 (1.597) | 0.034 (1.536) |
| CC_1/2_ | 0.952 (0.316) | 0.997 (0.451) |
| *<I>* /< σ*I>* | 4.0 (0.2) | 11.6 (0.2) |
| Completeness (%) | 100 (100) | 100 (100) |
| Redundancy | 51.6 (54.4) | 78.7 (78.8) |
| **Refinement** | |  |
| Resolution (Å) | 35.0–4.36 | 55.3–2.42 |
| No. reflections | 17940 | 94155 |
| *R*_work_ /*R*_free_ | 0.331/0.315 | 0.213/0.239 |
| No. atoms | 4861 | 10072 |
| Average *B*-factors ( Å^2^) | 151 | 89 |
| Parameters |  |  |
| Positional | 4298 | N/A |
| Flexibility | 2391 | N/A |
| Total | 6689 | N/A |
| R.m.s. deviations |  |  |
| Bond lengths (Å) | N/A | 0.002 |
| Bond angles (**°**) | N/A | 0.5 |

Numbers in brackets refer to the highest resolution shell of data

**Table S4** **| Cryo-EM data collection parameters**

|  | **3h incubation** | **50 min incubation**  **trimer** | **50 min incubation**  **‘dimer’** |
| --- | --- | --- | --- |
| **Data collection and reconstruction** | | | |
| Voltage (kV) | 300 | | |
| Frames | 40 | 40 | |
| Dose rate (e^-^/ Å ^2^/ s) | 20.2 | 20.7 | |
| Total dose (e^-^/ Å ^2^) | 42 | 42.0 | |
| Pixel size (Å) (super-resolution) | 0.415 | 0.415 | |
| Defocus (μm) | 0.8-2.6 | 0.8-2.6 | |
| Symmetry | C1 | C1 | |
| Movies | 7,032 | 13,307 | |
| Particles | 24,303 | 327,945 | 100,295 |
| Resolution FSC = 0.143 (Å) |  | 3.3 | 3.9/7.0 |
| Map sharpening B-factor (Å^2^) |  | -111.1 | -92.3 |
| **Model refinement** | | | |
| Model-to-map fit, CC_mask |  | 0.84 | 0.47 |
| R.m.s.d., bonds (Å) |  | 0.006 | 0.002 |
| R.m.s.d., angles (°) |  | 0.9 | 0.5 |
| All-atom Clash score |  | 7.6 | 1.8 |
| Rotamer outliers (%) |  | 3.8 | 2.0 |
| Ramachandran plot | | | |
| Favored (%) |  | 95.4 | 95.9 |
| Allowed (%) |  | 3.8 | 3.9 |
| Outliers (%) |  | 0.2 | 0.2 |
